# Antibody-lectin chimeras for glyco-immune checkpoint blockade

**DOI:** 10.1101/2022.10.26.513931

**Authors:** Jessica C. Stark, Melissa A. Gray, Simon Wisnovsky, Itziar Ibarlucea-Benitez, Marta Lustig, Nicholas M. Riley, Mikaela K. Ribi, Wesley J. Errington, Bence Bruncsics, Casim A. Sarkar, Thomas Valerius, Jeffrey V. Ravetch, Carolyn R. Bertozzi

**Affiliations:** Department of Chemistry and Sarafan ChEM-H, Stanford University, Stanford, CA, USA; Current affiliation: Department of Biological Engineering, Department of Chemical Engineering, and Koch Institute for Integrative Cancer Research, Massachusetts Institute of Technology, Cambridge, MA, USA; Faculty of Pharmaceutical Sciences, University of British Columbia, Vancouver, BC, Canada; Laboratory of Molecular Genetics and Immunology, The Rockefeller University, New York, NY, USA; Division of Stem Cell Transplantation and Immunotherapy, Department of Medicine II, Christian Albrechts University Kiel and University Medical Center Schleswig-Holstein, Kiel, Germany; Department of Biomedical Engineering, University of Minnesota, Minneapolis, MN, USA; Department of Measurement and Information Systems, Budapest University of Technology and Economics, Budapest, Hungary; Howard Hughes Medical Institute, Stanford, CA, USA

## Abstract

Despite the curative potential of checkpoint blockade immunotherapy, most patients remain unresponsive to existing treatments. Glyco-immune checkpoints – interactions of cell-surface glycans with lectin, or glycan-binding, immunoreceptors – have emerged as prominent mechanisms of immune evasion and therapeutic resistance in cancer. Here, we describe antibody-lectin chimeras (AbLecs), a modular platform for glyco-immune checkpoint blockade. AbLecs are bispecific antibody-like molecules comprising a cell-targeting antibody domain and a lectin “decoy receptor” domain that directly binds glycans and blocks their ability to engage inhibitory lectin receptors. AbLecs potentiate anticancer immune responses including phagocytosis and cytotoxicity, outperforming most existing therapies and combinations tested. By targeting a distinct axis of immunological regulation, AbLecs synergize with blockade of established immune checkpoints. AbLecs can be readily designed to target numerous tumor and immune cell subsets as well as glyco-immune checkpoints, and therefore represent a new modality for cancer immunotherapy.

## Main

T cell checkpoint blockade immunotherapies (e.g., PD-1/PD-L1 or CTLA-4 targeting antibodies) are now used to treat nearly half of all cancer patients in the US^1^. However, these therapies are effective in only a small fraction (<20%) of treated patients across cancer types^1^ and, of those who respond, at least 25% will develop resistance^2^. Thus, there is an urgent need for therapies targeting additional immune checkpoints that drive cancer progression.

Altered cell surface glycosylation is a hallmark of cancer that is associated with poor disease prognosis^3–5^. These altered glycan structures modulate immune cell function through interactions with lectin (glycan-binding) immunoreceptors, including sialic acid-binding immunoglobulin-like lectins (Siglecs)^6^, selectins^7^, and galectins^8,9^. In particular, multiple recent studies indicate that upregulation of cell-surface sialoglycans allows tumors to engage inhibitory Siglec receptors on immune cells and evade immune surveillance^10–27^. Taken together, glyco-immune checkpoints are emerging as attractive targets for cancer immunotherapy.

However, multiple challenges have historically prevented therapeutic targeting of glyco-immune checkpoints in cancer (**Fig 1a**). The weak immunogenicity of mammalian glycan structures can undermine the development of specific and high-affinity anti-glycan antibodies. To put this in perspective, a recent report cataloged 417 commercially available antibodies targeting glycans^28^. The same study noted that a single supplier offers 287 unique antibodies to tubulin and 269 unique antibodies to CD4, meaning that there are more antibodies to these two proteins available from a single supplier than all the commercially available anti-glycan antibodies combined. Further complicating monoclonal antibody development is the fact that many lectin receptors have multiple ligands that are expressed in the context of cancer^10,15,17,23,29^, and often the precise identities of their ligands are unknown. “Decoy receptor” molecules composed of a lectin glycan-binding domain fused to an antibody Fc domain address this complexity by retaining native glycan-binding specificities; however, their low binding affinities (typically with K_D_s in the μM to mM range^30^) preclude their use as therapeutics.

**Figure 1.**
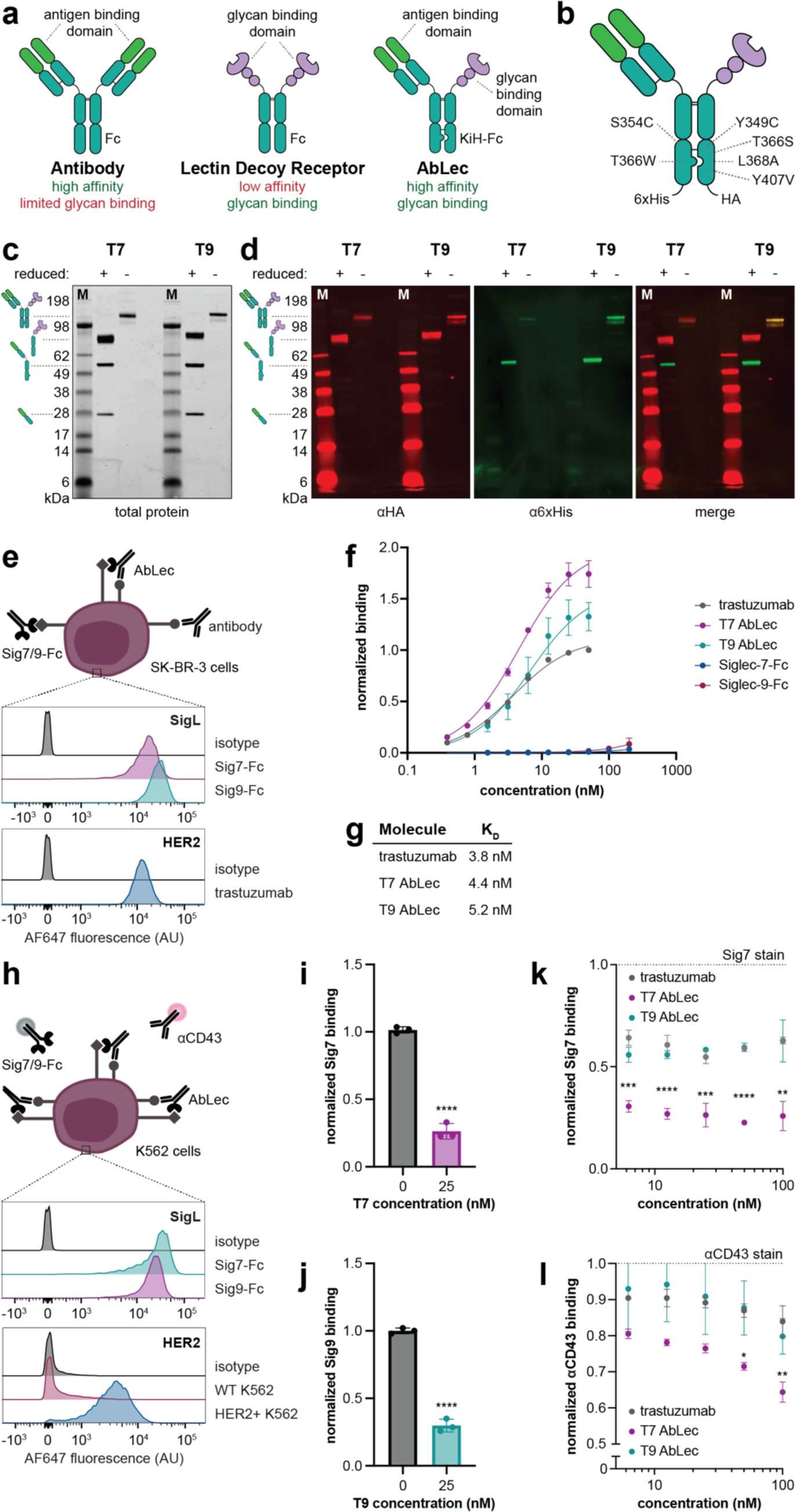
AbLecs enable the use of lectin decoy receptors for glyco-immune checkpoint blockade. (a) AbLecs couple tumor-targeting antibodies to glycan binding domains from lectin immunoreceptors, enabling blockade of immunosuppressive glycans at therapeutically relevant concentrations. (b) Fc engineering in a human IgG1 antibody framework facilitates self-assembly of AbLecs. (c) Total protein stain of SDS-PAGE of purified T7 and T9 AbLecs under disulfide-reducing and non-reducing conditions. Gel is representative of n = 3 replicates. (d) Western blot detection of antibody heavy chain (α6xHis) and Siglec-Fc chains (αHA) revealed that AbLecs are composed of antibody and Siglec decoy receptor arms. Blots are representative of n = 3 replicates. (e) SK-BR-3 cells express the HER2 antigen bound by trastuzumab as well as ligands for Siglec-7 and -9 (Sig7L and Sig9L, respectively) as measured by Siglec-Fc binding and analyzed via flow cytometry. Histograms are representative of n = 3 biological replicates (quantified in **Extended Data Fig. 2c-d**). (f) Binding of T7 AbLec, T9 AbLec, trastuzumab, Siglec-7-Fc, and Siglec-9-Fc to SK-BR-3 cells was quantified via flow cytometry. Data are mean ± standard deviations (s.d.) of n = 3 biological replicates, normalized to staining with 50 nM trastuzumab for each replicate. (g) Dissociation constants for trastuzumab, T7, and T9 AbLecs determined by fitting experimental data from (**f**) to a one-site total binding curve. (h) HER2+ K562 cells express the HER2 antigen as well as Sig7L and Sig9L via flow cytometry. Histograms are representative of n = 3 biological replicates (quantified in **Extended Data Fig**. **2c-d)**. (**i, j**) T7 (**i**) or T9 (**j**) AbLec treatment blocks Siglec-7 or Siglec-9 receptor binding, respectively, in competitive binding assays with dye labeled Siglec-7/9-Fc. Data are mean ± s.d. of n = 3 biological replicates, normalized to Siglec-Fc staining without AbLec. (k) T7 AbLec, but not trastuzumab or T9 AbLec, blocks Siglec-7 receptor binding in competitive binding assays with dye labeled Siglec-7-Fc. Data are mean ± s.d. of n = 3 biological replicates, normalized to Siglec-7-Fc staining with 0 nM AbLec or antibody. (l) Detection of αCD43 antibody (MEM-59) binding in competitive binding assays with increasing concentrations of T7 AbLec, T9 Ablec, or trastuzumab. The MEM-59 antibody binds to the same epitope on CD43 recognized by the Siglec-7 receptor. Data are mean ± s.d. of n = 3 biological replicates, normalized to αCD43 staining without AbLec or antibody.

We recently developed enzyme conjugates for targeted degradation of glycans^12,16^ or glycoproteins^31^ as an alternative approach to potentiate anti-tumor immunity, but these molecules require substantial engineering of enzyme kinetics in order to target new types of glycans. In addition, such approaches are currently restricted to degradation of broad classes of glycans (e.g., sialoglycans^12,16^) or glycoproteins (e.g., mucins^31^), rather than the specific molecules involved in immune regulation. Finally, while lectin-targeted blockade antibodies such as those directed against galectin-9^32^ and Siglec-15^33^ as well as a sialidase-Fc fusion for sialoglycan degradation^34^ are being tested in the clinic, these untargeted checkpoint blockade reagents could elicit immune-related adverse events, which are observed in at least 50% of patients treated with existing checkpoint blockade immunotherapies^35^. In contrast, targeted immunotherapies (e.g., monoclonal antibodies targeting tumor-associated antigens) generally offer more favorable safety profiles^36^. Overall, our inability to specifically and directly block immunomodulatory glycans, even after development of multiple therapeutic modalities, highlights the significance of this technological gap.

Here, we describe antibody-lectin (AbLec) chimeras as a modular platform to target glycans for cancer immunotherapy. AbLecs couple glycan binding domains from lectin receptors to antibodies directed against an antigen on the target cell surface (**Fig. 1a**). This approach uniquely enables blockade of immunomodulatory glycans at nanomolar concentrations agnostic of their specific identities. We demonstrate proof-of-concept of the AbLec platform through the development and characterization of AbLecs combining tumor-targeting antibodies with decoy receptor domains from Siglecs-7 and -9, which have been shown to mediate immune suppression in multiple cancers^11–13,16–27^. We show that tumor-targeting AbLecs enhance antibody-dependent phagocytosis and cytotoxicity of cancer cells by multiple primary immune cell subsets compared to the parent monoclonal antibody. AbLecs outperformed combination of a tumor-targeting antibody and a lectin receptor-blocking antibody, suggesting there is a therapeutic benefit gained by targeting glycans directly. We show that the AbLec platform can be applied to target diverse tumor-associated antigens, including HER2, CD20, and EGFR, and cancer cells with varying levels of antigen expression. Further, we designed and validated dual blockade AbLecs that simultaneously block protein-based immune checkpoints (e.g., PD-1/PD-L1, CD47/SIRPα) as well as glyco-immune checkpoints known to play roles in cancer (e.g., Siglec-7^13,15,19,20,25^, galectin-9^37,38^). We found that glyco-immune checkpoint blockade was synergistic with blockade of these established checkpoints. These results indicate that AbLecs target a non-redundant axis of immune suppression in cancer and could expand the subset of patients who respond to immunotherapy. In sum, AbLecs represent a generalizable approach for glyco-immune checkpoint blockade and a new modality for cancer immunotherapy.

## Results

### AbLecs bind to targeted cells at nanomolar concentrations

We hypothesized that high-affinity binding of an antibody Fab domain to tumor antigens would allow AbLecs to accumulate at sufficiently high local concentrations to permit binding of a relatively low affinity lectin domain to inhibitory glycans on the same cell. As proof-of-concept, we created AbLecs that combine the Fab domain from trastuzumab, an FDA-approved monoclonal antibody that binds the tumor antigen HER2, with the extracellular domains of Siglec-7 or -9 fused to an Fc domain. We used a modified knobs-into-holes Fc engineering approach^39^ to facilitate self-assembly of heterotrimeric AbLecs in a single expression culture (**Fig. 1b**). Indeed, coexpression of trastuzumab heavy and light chains with Siglec-7-Fc or Siglec-9-Fc in Expi293 cells resulted in expression of predominant protein species with molecular weights consistent with heterotrimeric AbLecs (**Fig. 1c**). Reducing SDS-PAGE analysis of purified proteins showed that trastuzumab x Siglec-7 (T7) and trastuzumab x Siglec-9 (T9) AbLecs were composed of 3 disulfide bonded protein chains consistent with the molecular weights of the Siglec-Fc and the trastuzumab heavy and light chains (**Fig. 1c**). Western blotting against HA and 6xHis tags on the Siglec-Fc and antibody heavy chains, respectively, further demonstrated that full-length AbLecs are composed of both antibody and decoy receptor arms (**Fig. 1d**), which was confirmed by mass spectrometry (**Extended Data Fig. 1**).

We next characterized binding of T7 and T9 AbLecs to HER2+ cell lines compared to trastuzumab and Siglec-Fc decoy receptor controls. Dissociation constant (K_D_) values for T7 and T9 AbLecs were measured by quantifying binding to SK-BR-3 cells at various concentrations via flow cytometry. SK-BR-3 cells express the HER2 antigen bound by trastuzumab as well as ligands for Siglecs-7 and -9 (Sig7/9L) (**Fig. 1e**). As expected, the decoy receptor controls (Siglec-7/9-Fc) bind only at low levels to SK-BR-3 cells, even at the highest concentrations tested (200 nM), despite the fact that SK-BR-3 cells express Siglec-7 and -9 ligands (**Fig. 1f**). However, by combining the decoy receptor arm with the high affinity trastuzumab antibody arm, AbLecs bind to SK-BR-3 cells with K_D_ values in the low nM range, similar to that of the parent bivalent antibody trastuzumab (**Fig. 1f, g**). Thus, AbLecs enable recruitment of otherwise low affinity lectin binding domains to cell surfaces at nanomolar concentrations.

Despite the low affinity of the Siglec domain in comparison to the antibody arm, AbLec binding affinity is influenced by both components. By mutating a conserved arginine residue in the Siglec-Fc binding site to an alanine, we created T7A and T9A AbLec mutants that are reported to exhibit significantly reduced affinities for Siglec ligands^40^. Arginine mutant AbLecs exhibited reduced binding to SK-BR-3 cells, measured by a ∼2-3-fold increase in apparent K_D_ compared to WT AbLecs (**Extended Data Fig. 2a-c**).

We further interrogated the mechanism of AbLec binding via a structure-guided computational modeling approach^41^. We used the recently developed *MVsim* toolset^42^ to construct a computational model of T7 AbLec binding trained using our experimental data (**Extended Data Fig. 2d**). Our computational model estimated that the apparent concentration of Siglec-7 ligands on the cancer cell surface was >2 orders of magnitude higher than that of the HER2 antigen (**Extended Data Fig. 2e**; [HER2]_app_ = 0.2 μM; [Sig7L]_app_ = 27 μM), suggesting that the abundance of accessible glycan ligands offsets the low monovalent Siglec domain affinity and facilitates cooperative AbLec binding. This observed contribution of the Siglec-Fc arm to binding may explain why AbLec dissociation constants are of the same order of magnitude as the bivalent parent antibody trastuzumab, despite having only one antibody arm.

### AbLecs selectively block lectin receptor binding epitopes

We next asked whether AbLecs could successfully block engagement of their targeted glycan ligands by competing with lectin receptors. Toward this end, we tested the ability of T7 and T9 AbLecs to compete with fluorescently-labeled Siglec decoy receptors for binding to HER2+ K562 cells. This cell line expresses a higher ratio of Sig7L and Sig9L to the targeted antigen HER2, offering a more challenging target for glycan blockade (**Fig. 1h**). Both T7 and T9 AbLecs bound to HER2+ K562 cells with low nanomolar K_D_ values (**Extended Data Fig. 2f, g**). Treatment of cells with T7 or T9 AbLec significantly reduced binding of fluorescently labeled Siglec-7 or Siglec-9, respectively, by flow cytometry (**Fig. 1i, j**). Importantly, AbLec binding to target cells is not cytotoxic: we did not observe any defects in SK-BR-3 or HER2+ K562 cell growth over 72 hours following AbLec treatment (**Extended Data Fig. 2h-k**). AbLecs further selectively block binding of the targeted lectin. Treatment with the T7 AbLec blocked binding of the cognate Siglec-7 decoy receptor to a greater extent than trastuzumab or the non-cognate T9 AbLec at all concentrations tested (**Fig. 1k**).

We wanted to test whether AbLecs block the same epitopes bound by their cognate lectin receptors. We recently reported that the predominant ligand for Siglec-7 expressed on K562 cells is the sialomucin CD43^17^. Further, we found that the commercially available MEM-59 antibody binds to the same sialylated epitope on CD43 that is bound by Siglec-7^17^. We tested the ability of AbLecs to compete with the MEM-59 antibody for binding to HER2+ K562 cells. We observed that the T7 AbLec blocked binding of MEM-59 to a greater extent than trastuzumab or the non-cognate T9 AbLec, suggesting that the T7 AbLec binds and blocks the same CD43 glycoform bound by the endogenous Siglec-7 immunoreceptor (**Fig. 1l**). Taken together, this evidence suggests that AbLecs selectively block engagement of the targeted lectin immunoreceptor by competing for the same binding epitopes.

### AbLecs potentiate antibody-dependent phagocytosis and cytotoxicity of cancer cells

Having demonstrated that AbLecs block binding of Siglec receptors to target cells, we investigated whether AbLec treatment could enhance antibody-mediated cancer cell killing. Therapeutic tumor-targeting antibodies create immune synapses between antigens on the cancer cell surface and Fc receptors (FcRs) on immune cells. Engagement of FcRs activates the immune cell to perform antibody effector functions including antibody-dependent cellular phagocytosis (ADCP) and antibody-dependent cellular cytotoxicity (ADCC), which result in destruction of the targeted cancer cell. However, Siglec ligands on the cancer cell surface recruit inhibitory Siglec receptors to the immune synapse, dampening immune cell activation and restraining antibody effector functions^12,16,43^. In contrast, we hypothesized that AbLecs would block Siglec engagement at the immune synapse and enhance antibody effector functions (**Fig. 2a-b**).

**Figure 2.**
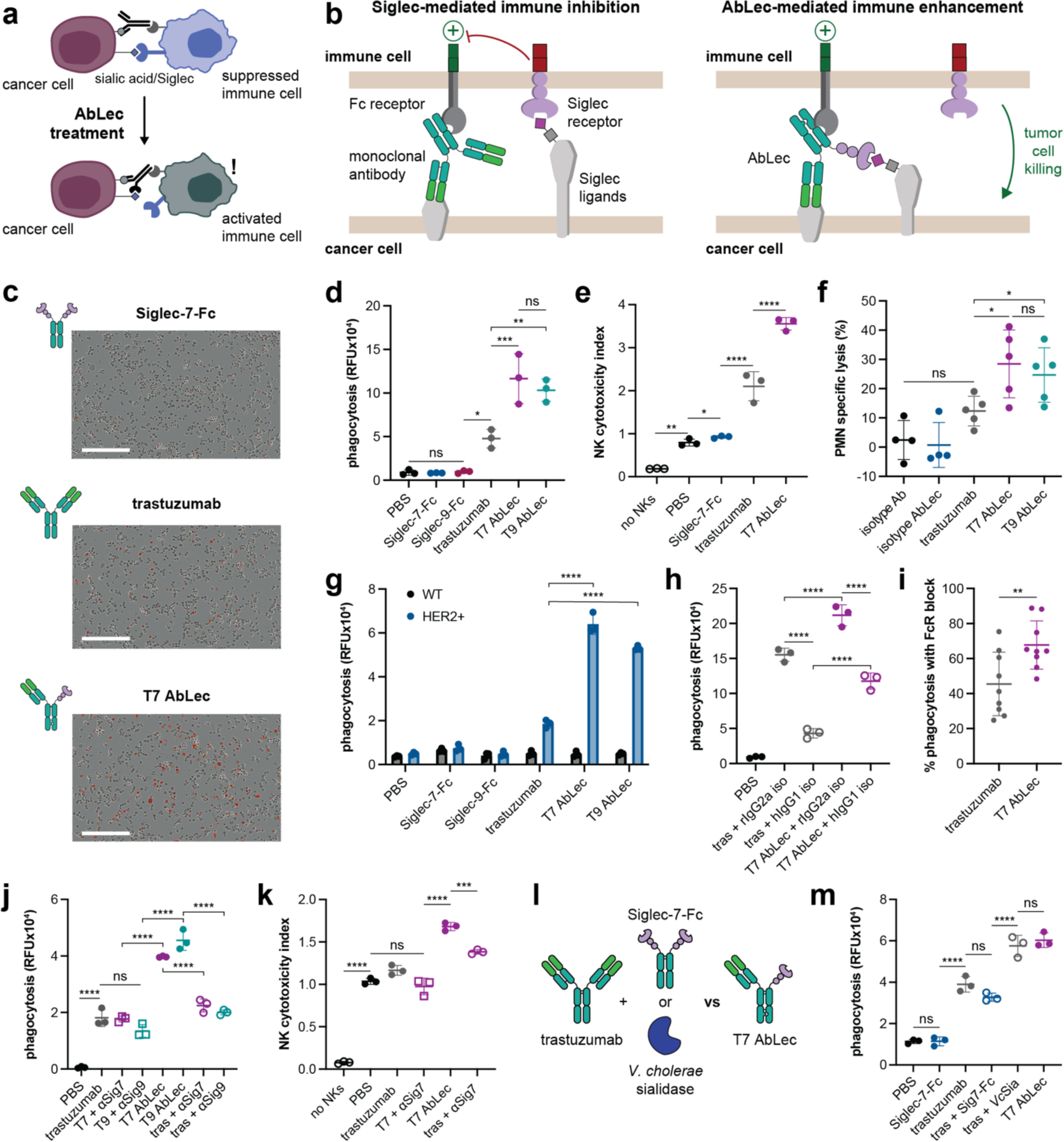
AbLecs enhance antibody effector functions via glyco-immune checkpoint blockade. (a) Siglec engagement at immune synapses restrains anti-tumor immune responses. We hypothesize that AbLecs will recruit FcR+ immune cells to tumors and simultaneously block Siglec engagement, relieving inhibitory signaling and potentiating cancer cell killing. (b) The chimeric AbLec architecture is designed to simultaneously elicit antibody effector functions via FcR engagement and block binding of glyco-immune checkpoint receptors. (c) Phase contrast and fluorescence microscopy images of primary human macrophage/SK-BR-3 cell co-cultures treated with Siglec-7-Fc, trastuzumab, or T7 AbLec at t = 5 h. SK-BR-3 cells were stained with the pHrodo red pH sensitive dye which fluoresces upon SK-BR-3 phagocytosis by macrophages. Images are representative of n = 3 biological replicates, scale bar represents 400 μm. (d) Phagocytosis of SK-BR-3 cells by primary human macrophages treated with T7/9 AbLecs, trastuzumab, or Siglec-7/9-Fc. Data are mean ± s.d. of n = 3 biological replicates. (e) Cytotoxicity of SK-BR-3 cells elicited by primary human NK cells treated with T7 AbLec, trastuzumab, or Siglec-7-Fc. Data are mean ± s.d. of n = 3 biological replicates. (f) Cytotoxicity of SK-BR-3 cells elicited by primary human granulocytes (PMNs) treated with T7/9 AbLecs, trastuzumab, or Siglec-7/9-Fc. Data are mean ± s.d. of biological replicates from n = 4-5 donors. (g) Phagocytosis of HER2+ K562 cells compared to WT K562 cells, which are HER2-, (Fig. 1h**; Extended Data Fig. 4c**) treated with T7/9 AbLecs, trastuzumab, or Siglec-7/9-Fc. Data are mean ± s.d. of n = 3 biological replicates. (h) Primary human macrophages were pre-treated with non-targeting antibodies to block human FcRs (hIgG1 iso) or non-FcR binding controls (rIgG2a iso) prior to co-culture with SK-BR-3 cells and treatment with trastuzumab (tras) or T7 AbLec. Data are mean ± s.d. of n = 3 biological replicates. (i) Percent phagocytosis observed upon FcR blockade (+ hIgG1 iso RFU/+ rIgG2a iso RFU). Data are mean ± s.d. of n = 3 biological replicates from each of n = 3 donors. (j) Macrophage ADCP of SK-BR-3 cells induced by trastuzumab or T7/9 AbLecs in the presence or absence of Siglec-7/9 blocking antibodies (αSig7 or αSig9, respectively). Data are mean ± s.d. of n = 3 biological replicates. (k) NK cell ADCC of SK-BR-3 cells induced by trastuzumab or T7 AbLec in the presence or absence of a Siglec-7 antagonist antibody (αSig7). Data are mean ± s.d. of n = 3 biological replicates. (l) We compared combination of trastuzumab with Siglec-7-Fc or *V. cholerae* sialidase (*Vc*Sia) to T7 AbLec treatment in their ability to activate macrophage ADCP. (m) Macrophage ADCP of HER2+ K562 cells treated with Siglec-7-Fc, trastuzumab, T7 AbLec, or combinations of trastuzumab with Siglec-7-Fc or *V. cholerae* sialidase (*Vc*Sia). Data are mean ± s.d. of n = 3 biological replicates.

We first tested our hypothesis using *in vitro* phagocytosis assays with primary human macrophages (**Extended Data Fig. 3a, b**). At the time of use in our assays, macrophages primarily expressed FcγRI and FcγRIIa (**Extended Data Fig. 3c, d**), as well as Siglecs-7 and -9 (**Extended Data Fig. 3e, f**). We co-cultured these macrophages with SK-BR-3 cells labeled with a pH-sensitive dye that fluoresces red in acidic phagosomes (**Extended Data Fig. 3a**), enabling quantification of phagocytosis via fluorescence. After 5 h of co-culture, fluorescence microscopy images show that SK-BR-3 cells in co-cultures treated with the T7 AbLec are more readily phagocytosed by macrophages compared to those treated with trastuzumab or Siglec-7-Fc decoy receptor (**Fig. 2c**). We quantified the observed increase in phagocytosis using macrophages from three unique donors and found that both T7 and T9 AbLecs significantly enhance phagocytosis of tumor cells compared to trastuzumab or Siglec-Fc decoy receptors alone (**Fig. 2d; Extended Data Fig. 4a, b**). We observed the same enhancement of phagocytosis of the HCC-1854 and HER2+ K562 cell lines, which express varying levels of HER2 and/or Sig7/9L compared to SK-BR-3 cells (**Extended Data Fig. 4c-j**).

We next asked whether AbLecs could potentiate cancer cell killing by additional subsets of immune cells. We co-cultured primary human NK cells expressing FcγRIII and Siglec-7 with SK-BR-3 tumor cells and analyzed cytotoxicity via flow cytometry (**Extended Data Fig. 5a-g**). Across three donors, the T7 AbLec significantly enhanced NK cell cytotoxicity compared to trastuzumab or Siglec-7-Fc decoy receptor (**Fig. 2e; Extended Data Fig. 5h, i**). We observed the same effect in assays with the HER2+ K562 cell line (**Extended Data Fig. 5j, k**). We performed similar assays with polymorphonuclear granulocytes (PMNs), which express FcγRIIa and FcγRIII^26^ as well as Siglecs-7 and -9 (**Extended Data Fig. 6a-d**). We found that T7 and T9 AbLecs elicited increased tumor killing by PMNs compared to trastuzumab and AbLec isotype controls (**Fig. 2f; Extended Data Fig. 6e-h**).

It was important to demonstrate that AbLec activity is at least partially antibody-dependent; that is, mediated via antigen binding and FcR engagement. We observed that T7/9 AbLec-mediated phagocytosis of target cells was dependent on expression of the HER2 antigen in phagocytosis assays with WT (HER2-) and HER2+ K562 cells (**Fig. 2g**). To test FcR dependence, we incubated macrophages with non-targeting antibodies of either rat IgG2a (rIgG2a) or human IgG1 (hIgG1) isotype prior to co-culture with SK-BR-3 cells. The human IgG1 antibody competes with both trastuzumab and T7 AbLec, which have hIgG1 Fc domains, to bind to macrophage FcRs. In contrast, the rIgG2a isotype has very low affinity for human FcRs^44^, and serves as a negative control for FcR blockade. We observed lower levels of SK-BR-3 phagocytosis for both trastuzumab and T7 AbLec when macrophages were pre-treated with hIgG1 isotype antibodies, indicating that both antibody and AbLec activity is FcR-dependent (**Fig. 2h; Extended Data Fig. 7a**). We further found that an antibody blocking FcγRIII/CD16 significantly reduced AbLec-mediated ADCC by NK cells, confirming that AbLec activity is FcR-dependent (**Extended Data Fig. 7b**). Notably, FcR blockade appeared to reduce trastuzumab-mediated phagocytosis to a greater extent than the T7 AbLec. Indeed, when we quantified the reduction in phagocytosis observed upon FcR blockade across three unique donors we found that trastuzumab exhibited a greater dependence on FcRs than the T7 AbLec (**Fig. 2i**). These data support our hypothesis that AbLec mechanism of action is bifunctional, with contributions from activation of antibody effector functions and glyco-immune checkpoint blockade. Overall, these results show that AbLecs elicit enhanced antibody effector functions including ADCP and ADCC compared to their parent monoclonal antibodies.

### AbLec efficacy is mediated via glyco-immune checkpoint blockade

Critically, we tested whether AbLec-mediated enhancement of anti-tumor immune responses was a result of blockade of Siglec-sialoglycan interactions. In these experiments, we pre-incubated macrophages or NK cells with Siglec-7 or -9 blocking antibodies prior to co-culture with SK-BR-3 cells (**Extended Data Fig. 7c, top**). If the targeted Siglec receptor was blocked, AbLec-mediated ADCP (**Fig. 2j; Extended Data Fig. 7d**) and ADCC (**Fig. 2k; Extended Data Fig. 7e**) were reduced to similar levels as those observed with trastuzumab treatment. However, if Siglec receptors were not blocked, AbLec treatment enhanced ADCP (**Fig. 2j; Extended Data Fig. 7d**) and ADCC (**Fig. 2k; Extended Data Fig. 7e**) compared to trastuzumab. We observed the same evidence of Siglec dependence in ADCP and ADCC assays with HER2+ K562 cells (**Extended Data Fig. 7f, g**). Pretreatment of target cells with sialidase to degrade Siglec ligands also abolished AbLec-mediated enhancement of PMN ADCC over trastuzumab (**Extended Data Fig. 6f, g**), further supporting the idea that AbLec activity is Siglec-dependent.

We further assessed the Siglec-dependence of AbLec activity in assays with murine bone marrow-derived macrophages (BMDMs). We isolated BMDMs from WT mice and co-cultured them with the syngeneic HER2+ B16D5 cell line, which expresses high levels of the HER2 antigen and Sig9L, and intermediate levels of Sig7L (**Extended Data Fig. 8a-c**). We measured activation of BMDMs via secretion of pro-inflammatory cytokines and chemokines upon treatment with AbLecs and controls. We did not observe significant levels of BMDM activation upon treatment with the T7 AbLec, perhaps due to the relatively low level of Sig7L expression on the HER2+ B16D5 cell line (**Extended Data Fig. 8a**). In contrast, T9 AbLec treatment significantly increased BMDM activation (**Extended Data Fig. 8d-g**). We established similar models using BMDMs from mice with engineered Siglec expression^20^. These included mice lacking Siglec-E, the murine ortholog for human Siglec-7/9, and mice humanized for Sig7/9, in which SigE was knocked out and human Sig7/9 receptors were knocked in (**Extended Data Fig. 8h**). In BMDMs from humanized Sig7/9 mice, we observed an increase in BMDM activation in co-cultures treated with T9 AbLec compared to those treated with trastuzumab. However, in SigE knockout mice, overall activation was increased and activation upon T9 AbLec treatment was indistinguishable from treatment with trastuzumab (**Extended Data Fig. 8i, j**). Taken together, these results suggest that the increased levels of ADCP and ADCC observed with AbLec treatment are a direct result of glycan blockade.

### The chimeric AbLec architecture enables potent glyco-immune checkpoint antagonism

Combination of monospecific therapies results in enhanced therapeutic efficacy, and is now thought to be required for durable responses in most cancer patients^45^. We hoped the bispecific AbLec architecture would elicit tumor-targeted antibody effector functions and glyco-immune checkpoint blockade at least as effectively as combination therapy. Thus, we sought to benchmark efficacy of AbLecs compared to combinations of trastuzumab with existing Siglec blockade approaches. First, we compared AbLec treatment to combinations of trastuzumab with Siglec blocking antibodies^20^ (**Extended Data Fig. 7c, bottom**). Surprisingly, AbLecs elicited enhanced ADCP (**Fig. 2j**) and ADCC (**Fig. 2k**) of SK-BR-3 cells compared to the combination of trastuzumab with Siglec-7 or -9 antagonist antibodies^20^. We observed the same trends in ADCP and ADCC assays with HER2+ K562 cells (**Extended Data Fig. 7f, g**). These data suggest that blockade of Siglec engagement locally, at the immune synapse, could be more effective than systemic Siglec antagonism.

We further compared AbLec efficacy to combination of trastuzumab with *Vibrio cholerae* sialidase as a model of enzymatic sialoglycan degradation, an approach to target the Siglec/sialic acid axis that is currently being investigated in clinical trials^34^ (**Fig. 2l**). The *Vibrio cholerae* sialidase cleaves all linkages (α2,3; α2,6; and α2,8) of terminal sialic acid, blocking Siglec engagement via ligand degradation^12,16^. There was no significant difference in ADCP elicited by T7 AbLec treatment compared to the combination of trastuzumab with *V. cholerae* sialidase (**Fig. 2l; Extended Data Fig. 9a, b**). Combination of T7 and T9 AbLecs did not further enhance ADCP compared to T7 or T9 treatment alone (**Extended Data Fig. 9c, d**). Similarly, activation of murine BMDMs treated with T9 AbLec was indistinguishable from combination treatment with trastuzumab with *V. cholerae* sialidase (**Extended Data Fig. 8d-g**). These data indicate that AbLec-mediated Siglec blockade is as effective as sialic acid degradation in its ability to potentiate antibody effector functions.

We asked whether the chimeric AbLec architecture was required to potentiate tumor killing. To do this, we compared phagocytosis in macrophage/HER2+ K562 cell cultures treated with T7 AbLec or the combination of trastuzumab with the Siglec-7-Fc decoy receptor (**Fig. 2l**). We found that combination of trastuzumab with Siglec-7-Fc did not enhance killing of HER2+ K562 cells compared to trastuzumab alone, as expected given the low binding affinity of the decoy receptor (**Fig. 2m; Extended Data Fig. 9a, b**). However, as before, treatment with T7 AbLec elicited enhanced tumor phagocytosis (**Fig. 2m; Extended Data Fig. 9a, b**). From these results, we conclude that the chimeric AbLec architecture is required to recruit lectin decoy receptor domains to tumor cell surfaces at therapeutically relevant concentrations and potentiate tumor killing.

Overall, our findings support the paradigm that the chimeric AbLec architecture results in a gain-of-function: the ability to block inhibitory signaling through glyco-immune checkpoints. AbLecs are more effective glyco-immune checkpoint antagonists than Siglec-blocking antibodies, and as effective as enzymatic sialic acid degradation, both of which are therapeutic strategies being investigated in clinical trials^33,34^.

### The AbLec platform is extensible to multiple cancers and mechanisms of action

Due to their modular chimeric architecture, AbLecs can be readily developed to target diverse cancers and glyco-immune checkpoints. To demonstrate this modularity, we constructed tumor-targeting AbLecs with components derived from rituximab x Siglec-7 (R7) and cetuximab x Siglec-7 (C7), designed to block Sig7L on CD20+ and EGFR+ cancer cells, respectively (**Fig. 3a, b**). In addition, we generated a trastuzumab x Siglec-10 AbLec that simultaneously targets HER2+ tumors while blocking engagement of the Siglec-10 receptor, which was recently shown to mediate immune evasion in breast and ovarian cancers^15^ (**Fig. 3c**).

**Figure 3.**
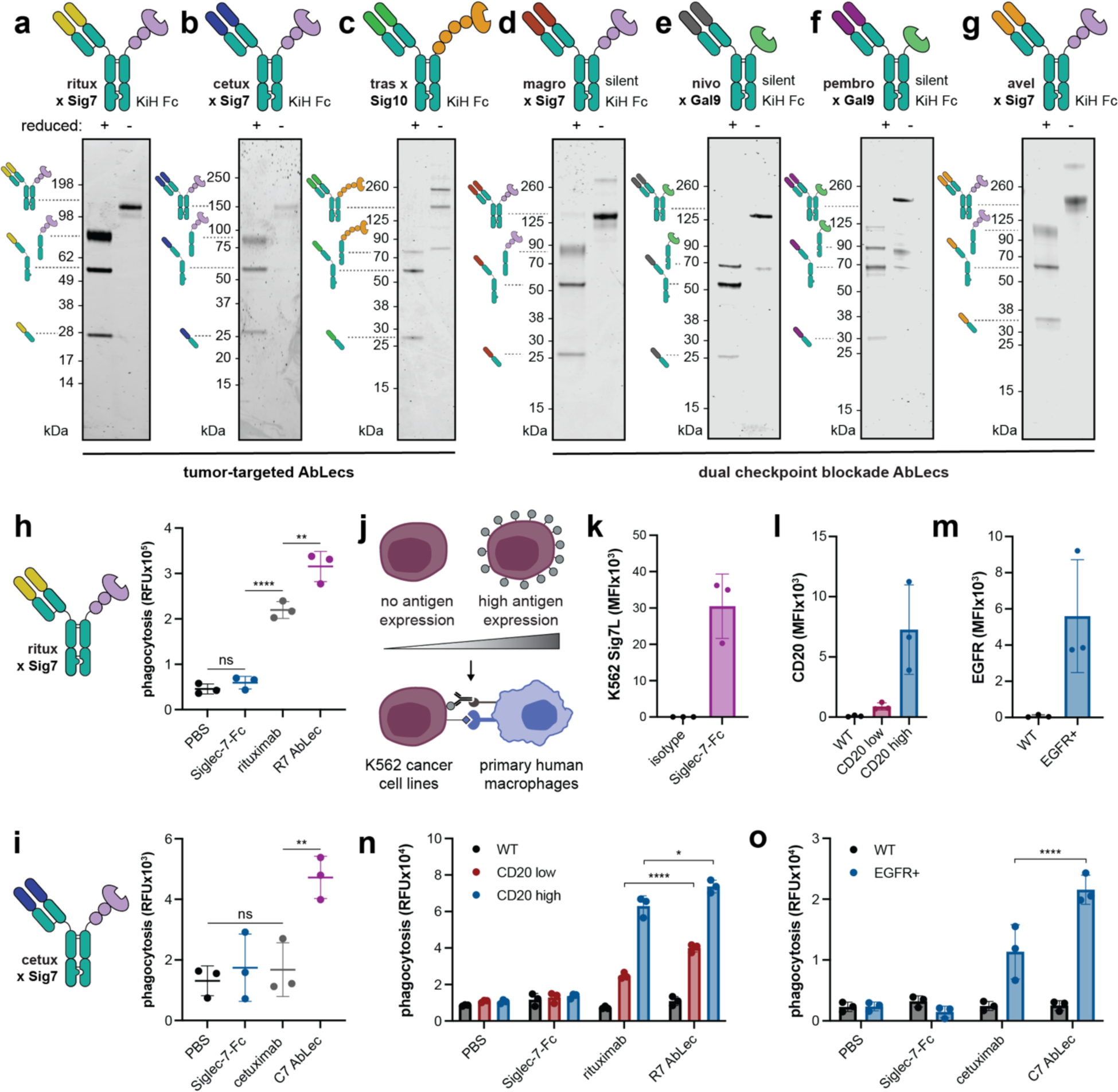
The AbLec platform is extensible to multiple cancers and mechanisms of action. (**a-g**) Reducing (+) and non-reducing (-) SDS-PAGE analysis of purified (**a**) rituximab (αCD20) x Siglec-7 (R7) AbLec, (**b**) cetuximab (αEGFR) x Siglec-7 (C7) AbLec, (**c**) trastuzumab (αHER2) x Siglec-10 (T10) AbLec, (**d**) magrolimab (αCD47) x Siglec-7 (M7) AbLec, (**e**) nivolumab (αPD-1) x Galectin-9 (NG9) AbLec, (f) pembrolizumab (αPD-1) x Galectin-9 (PG9) AbLec, and (**g**) avelumab (αPD-L1) x Siglec-7 (A7) AbLec. Gels are representative of n = 3 replicates. (h) ADCP of Ramos cells by primary human macrophages in co-cultures treated with R7 AbLec, rituximab, or Siglec-7-Fc. Data are mean ± s.d. of n = 3 biological replicates. (i) ADCP of EGFR+ K562 cells by primary human macrophages in co-cultures treated with C7 AbLec, cetuximab, or Siglec-7-Fc. Data are mean ± s.d. of n = 3 biological replicates. (j) We engineered K562 cell lines expressing high and low levels of tumor-associated antigens (CD20, EGFR). WT K562s were antigen negative. These cell lines were used to interrogate AbLec efficacy in tumors with low antigen density. (k) Sig7L expression measured via Siglec-7-Fc staining of K562 cells compared to hIgG1 isotype control. Data are mean ± s.d. of n = 3 biological replicates. (l) CD20 expression on WT, CD20 low, and CD20 high K562 cells measured via rituximab staining compared to hIgG1 isotype antibody by flow cytometry. Data are mean ± s.d. of n = 3 biological replicates. (m) EGFR expression on WT and EGFR+ K562 cells measured via cetuximab staining compared to hIgG1 isotype antibody by flow cytometry. Data are mean ± s.d. of n = 3 biological replicates. (n) Macrophage ADCP of WT, CD20 low, and CD20 high K562 cells treated with R7 AbLec, rituximab, or Siglec-7-Fc. Data are mean ± s.d. of n = 3 biological replicates. (o) Macrophage ADCP of WT and EGFR+ K562 cells treated with C7 AbLec, cetuximab, or Siglec-7-Fc. Data are mean ± s.d. of n = 3 biological replicates.

We further developed dual checkpoint blockade AbLecs designed to block engagement of both protein and glycan-based checkpoint receptors. We designed an AbLec for dual blockade of the “don’t eat me” signal CD47 and Siglec-7 ligands on tumor cells by combining the clinical-stage αCD47 antibody magrolimab^46^ with a Siglec-7 decoy receptor domain (**Fig. 3d**). We generated nivolumab x galectin-9 and pembrolizumab x galectin-9 AbLecs designed to simultaneously block the PD-1/PD-L1 and TIM-3/galectin-9 T cell checkpoints (**Fig. 3e, f**). Nivolumab and pembrolizumab are PD-1 blocking antibodies approved to treat diverse cancers^47–55^. Notably, the galectin-9 lectin is a ligand for the TIM-3 checkpoint receptor, and binding of galectin-9 to TIM-3 contributes to CD8+ T cell exhaustion^38,56^. Both galectin-9 and TIM-3 antagonist antibodies are cancer immunotherapy approaches being investigated in clinical trials^32,57–59^. Finally, we used the binding domain from the FDA-approved αPD-L1 antibody avelumab^60–62^ to engineer an avelumab x Siglec-7 AbLec for dual blockade of PD-L1 and Sig7L expressed on tumor or antigen presenting cells (**Fig. 3g**).

Functional characterization of R7 and C7 AbLecs demonstrated the utility of the AbLec platform for combination tumor targeting and glyco-immune checkpoint blockade in diverse tumor types. Across n = 3 unique donors, R7 and C7 AbLecs significantly enhanced phagocytosis of CD20+ Ramos cells or EGFR+ K562 cells compared to the parent antibody or Siglec-7-Fc alone (**Fig. 3h, i; Extended Data Fig. 9e-j**). Importantly, and consistent with our findings with T7/9 AbLecs, R7 and C7 AbLecs selectively target antigen expressing cells and spare antigen negative cells (**Fig. 3j-o**). Thus, AbLecs can be readily redesigned to target a variety of cancer antigens.

Low or heterogeneous expression of tumor-associated antigens on cancer cell surfaces has been associated with resistance to immunotherapies including monoclonal antibodies^63,64^, immune checkpoint blockade^65^, and CAR-T cells^66,67^. We hypothesized that the enhanced effector functions elicited by AbLecs might be able to overcome weak FcR signaling potentiated by tumors with low antigen density. We created K562 lines expressing high and low levels of the CD20 antigen, while the expression of Sig7L was constant (**Fig. 3j-l**). In assays with primary human macrophages, we found that R7 AbLec treatment enhanced ADCP of tumor cells expressing very low levels of the targeted CD20 antigen compared to the parent monoclonal antibody, rituximab (**Fig. 3j-l, n**). Our results show that the improved efficacy of AbLecs compared to the parent monoclonal antibody is maintained in cancers with low antigen expression, a current area of unmet medical need.

### AbLecs synergize with blockade of established immune checkpoints

Siglecs and sialic acids are upregulated in patients that do not respond to PD-1 blockade^27^, suggesting that glyco-immune checkpoints are an orthogonal mechanism of immune evasion in the tumor microenvironment. If this is true, we would expect AbLecs to synergize with existing checkpoint blockade modalities. We tested this hypothesis by investigating whether trastuzumab hybrid AbLecs synergize with blockade of CD47, in light of recent results showing that CD47 blockade synergizes with trastuzumab^68^. If glyco-immune checkpoints are a distinct axis of immune suppression, we would expect the combination of trastuzumab hybrid AbLecs with CD47 blockade to further enhance ADCP of HER2+ cancer cells (**Fig. 4a**). Indeed, combination of a CD47 antagonist antibody and T7 AbLec elicited enhanced tumor killing compared to T7 AbLec treatment alone in 2 of 3 donors tested (**Fig. 4b; Extended Data Fig. 10a-c**). Across all 3 donors, combination of αCD47 and T7 AbLec outperformed the combination of αCD47 and trastuzumab (**Extended Data Fig. 10a-c**). Combination of αCD47 and T7 AbLec also enhanced PMN cytotoxicity over αCD47 and trastuzumab (**Extended Data Fig. 6h**).

**Figure 4.**
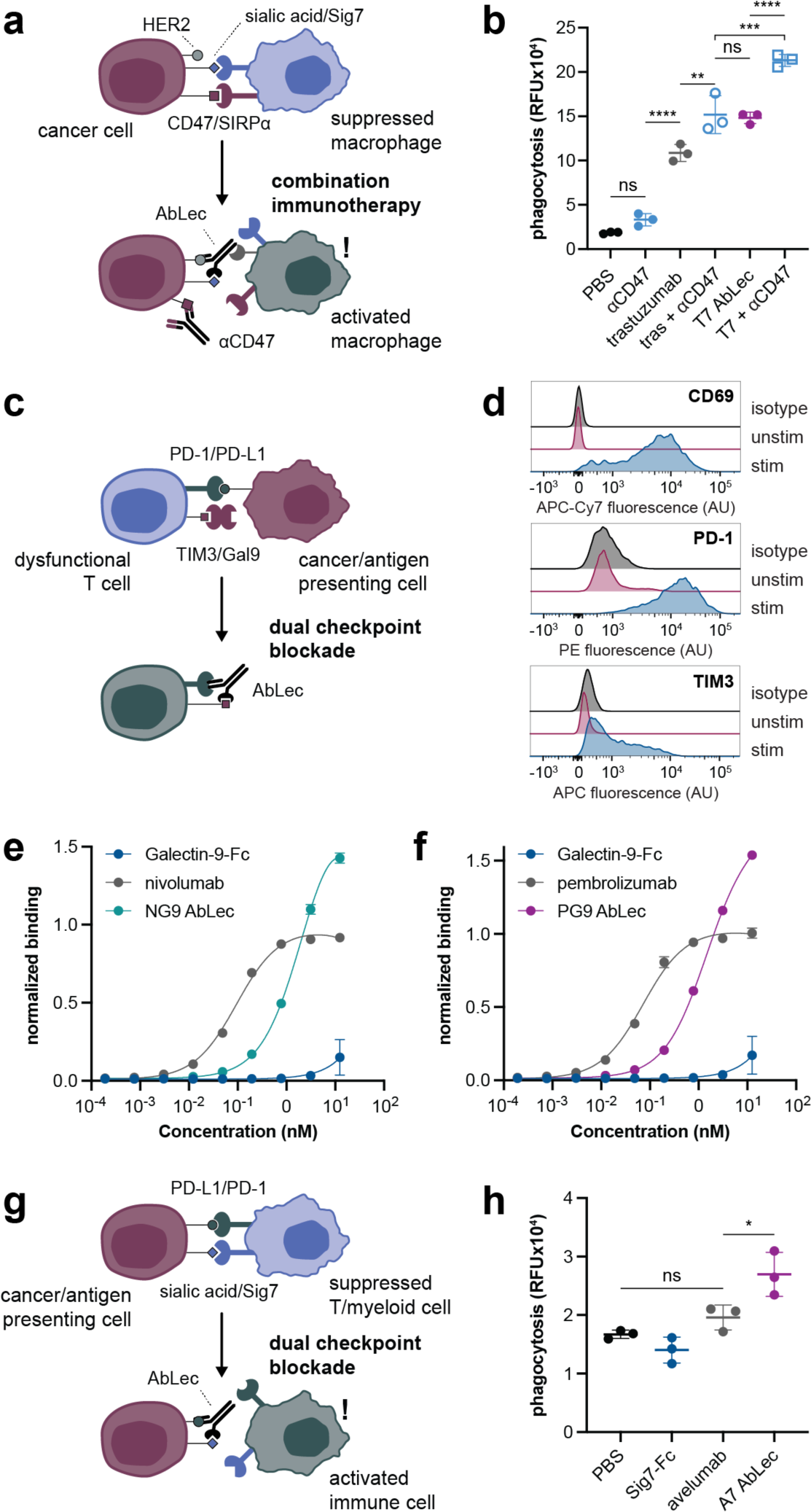
AbLecs target a distinct axis of immune regulation and can be used to simultaneously block protein- and glycan-based immune checkpoints. (a) We tested the ability of T7 AbLec treatment to synergize with CD47 blockade in ADCP assays with primary human macrophages. (b) ADCP of SK-BR-3 cells by primary human macrophages in co-cultures treated with trastuzumab or T7 AbLec alone and in combination with a CD47 antagonist antibody. Data are mean ± s.d. of n = 3 biological replicates. (c) We developed dual checkpoint blockade AbLecs to simultaneously block the PD-1/PD-L1 and TIM-3/galectin-9 T cell checkpoints. (d) Following 3 days of stimulation with αCD3/αCD28 antibodies, primary human CD3+ T cells upregulated CD69, PD-1, and TIM-3 by flow cytometry compared to unstimulated T cell and isotype controls. (**e-f**) NG9 (**e**) and PG9 (**f**) AbLecs bind to activated human CD3+ T cells at nanomolar concentrations. Data are mean ± s.d. of n = 3 biological replicates. (g) We developed dual checkpoint blockade AbLecs to simultaneously block the PD-1/PD-L1 and Siglec-7 checkpoints that negatively regulate activation of both T cells and myeloid cells. (h) Phagocytosis of MDA-MB-231 cells by primary human macrophages in co-cultures treated with A7 AbLec, avelumab (αPD-L1), or Siglec-7-Fc. Data are mean ± s.d. of n = 3 biological replicates.

In light of this finding, we designed dual checkpoint blockade AbLecs to enable blockade of protein- and glycan-based checkpoints with a single chimeric molecule. We first characterized AbLecs targeting the PD-1/PD-L1 and TIM-3/galectin-9 checkpoints on T cells, which combined the nivolumab or pembrolizumab αPD-1 antibody binding domains with the glycan-binding domain from galectin-9 (NG9 and PG9 AbLecs, respectively; **Fig. 4c**). We found that both AbLecs bound to activated human T cells at nanomolar concentrations (**Fig. 4d-f; Extended Data Fig. 10d**). Further, NG9 and PG9 AbLecs exhibited distinct binding profiles compared to parent monoclonal antibody and galectin-9-Fc controls (**Fig. 4e, f**), indicating contributions from both αPD-1 and galectin-9 domains to overall AbLec binding.

Finally, we asked whether dual checkpoint blockade AbLecs were functional (**Fig. 4g**). The A7 AbLec combines the Fab domain from the αPD-L1 antibody avelumab with the glycan-binding domain from Siglec-7 for dual blockade of the PD-1/PD-L1 and Siglec-7/sialic acid immunological axes. A7 AbLec enhanced phagocytosis of MDA-MB-231 cells expressing PD-L1 and Sig7L compared to the avelumab antibody alone (**Fig 4h; Extended Data Fig. 10e-g**). Taken together, AbLecs target an orthogonal immunosuppressive pathway in cancer and can be used alone or in combination with existing clinical-stage immunotherapies to drive enhanced antitumor immune responses.

## Discussion

In this work, we establish AbLecs as a new class of antibody chimeras targeting glyco-immune checkpoints for cancer immunotherapy. AbLecs combine antibodies targeting antigens of interest (e.g., tumor-associated antigens, immune checkpoint receptors or ligands) with glycan-binding domains from immunomodulatory lectin receptors. We show that AbLecs bind to cell surfaces at therapeutically relevant concentrations and selectively block binding of the targeted lectin receptor. Tumor-targeted AbLecs potentiated antibody effector functions including phagocytosis and cytotoxicity of cancer cells, including those with low expression of the targeted antigen. Synergy with blockade of established checkpoints demonstrated the potential for AbLecs to expand the subset of patients that benefit from cancer immunotherapy. Overall, AbLecs offer a modular molecular architecture for glycan blockade as a unique immunotherapeutic mechanism of action.

Unexpectedly, we found that T7 and T9 AbLecs were more effective Siglec antagonists than Siglec-blocking antibodies and as effective as sialoside degradation. This result suggests that proximity of the tumor-targeting and Siglec blockade elements may be important. This idea is consistent with a recent study demonstrating that Siglec recruitment to FcRs can fully inhibit FcR activation^43^. Similarly, emerging evidence indicates that recruitment of cell-surface phosphatases to immunoreceptors inhibits their activation^69^, while phosphatase exclusion potentiates signaling^70^. In light of these findings, we propose that the ability of tumor-targeted AbLecs to enhance immune cell activation is mediated via exclusion of inhibitory lectin receptors from the immunological synapse. Our data suggests that targeted lectin blockade mediated by AbLecs is at least as effective as systemic glyco-immune checkpoint blockade with the potential for reduced toxicity.

Tumors dramatically remodel their cell surface glycosylation. It is now clear that a key role for these remodeled glycans is to facilitate immune evasion and tumor progression by engaging multiple broad classes of lectin immunoreceptors^10–27,37,38,71–79^. Our chimeric AbLec molecules result in a gain-of-function: the ability to block glyco-immune checkpoints that represent an important mechanism of immune suppression and therapeutic resistance in cancer. The AbLec approach is extensible to multiple cancers and mechanisms of action, with the potential to expand the efficacy of cancer immunotherapy.

## Methods

### Cell lines

All cell lines were purchased from the American Type Culture Collection (ATCC) unless otherwise noted. SK-BR-3, HCC-1954, K562, and Ramos cells were cultured in RPMI + 10% fetal bovine serum (FBS) without antibiotic selection. HER2+ B16D5 cells were a gift from the Weiner laboratory (Lombardi Cancer Center) and were cultured in DMEM with 10% FBS, 1% glutamine, 1% sodium pyruvate and 1% amino acids and without antibiotic selection. MDA-MB-231 cells were cultured in DMEM with 10% FBS, 1% glutamine, 1% sodium pyruvate and 1% amino acids and without antibiotic selection. Expi293F cells were a gift from the Kim lab at Stanford and were cultured according to the manufacturer protocol (Thermo Fisher Scientific) protocol. K562s were transfected according to manufacturer’s protocol with EGFR using pre-packaged lentiviral particles (G&P Biosciences) and selected for EGFR expression by culture in 1 µg/mL puromycin (InVivoGen). K562s were transfected according to manufacturer’s protocol with CD20 using pre-packaged lentiviral particles (G&P Biosciences) and sorted for high and low CD20 expressing cells using rituximab and a BV421-labeled anti-human secondary (Jackson ImmunoResearch). Lentiviral vector encoding *HER2/neu* was a gift from Mien-Chie Hung (Addgene plasmid #16257). Stable HER2^+^ cell lines were generated as previously described^80^ and HER2 expression was verified by flow cytometry. Cell lines were not independently authenticated beyond the identity provided by the ATCC. Cell lines were cultivated in a humidified incubator at 5% CO_2_ and 37 °C and tested negative for mycoplasma quarterly using a PCR-based assay.

### Plasmids and protein sequences

All antibody and AbLec plasmids were generated by Twist Bioscience and inserted into the pTwist CMV BetaGlobin vector using the XhoI and NheI cut sites, unless otherwise specified below. DNA and amino acid sequences are listed in **Supplemental Table 1**. Trastuzumab was expressed from a pCDNA3.1 vector described in our previous work^16^. The rituximab and cetuximab antibody variable sequences were generated by IDT and cloned into the variable regions of the VRC01 antibody plasmid vector (a generous gift from the Kim lab at Stanford) by using the In-Fusion cloning kit (Takara) according to the manufacturer’s protocol. The EndoFree Plasmid Maxi Kit (Qiagen) was used to prepare DNA for transfection.

### Protein expression and purification

Antibodies and AbLecs were expressed via transient transfection in Expi293F cells (Thermo Fisher Scientific) according to the manufacturer protocol. For antibodies, a 1:1 heavy to light chain plasmid ratio by weight was used. The trastuzumab antibody heavy chain and light chain were co-expressed from a single plasmid. For AbLecs, a 2:1:1 ratio of lectin:heavy chain:light chain was used. After seven days of expression, proteins were collected from the supernatant by pelleting cells at 500 *x g* for 5 min, followed by clarification with a spin at 4000 *x g* for 40 min, and 0.2 μm filtration. Antibodies were purified by manual gravity column (BioRad) using protein A agarose (Fisher Scientific). Briefly, the clarified supernatant was flowed over the column twice, bound protein was sialidase treated on the resin with 0.25-0.5 column volumes of 100 nM *V. cholerae* sialidase in PBS for 0.5-2 h rt, washed with 20 column volumes of PBS, and eluted twice with 5 mL 100 mM glycine buffer pH 2.8 into tubes pre-equilibrated with 150 μL of 1 M Tris pH 8. AbLecs were purified by manual gravity column using nickel-NTA agarose resin (Qiagen). Briefly, AbLec supernatant was incubated with PBS-equilibrated resin for ∼1 hour at 4 °C. Resin and supernatant was then loaded onto a chromatography column, bound protein was sialidase treated on the resin with 0.25-0.5 column volumes of 100 nM *V. cholerae* sialidase in PBS for 0.5-2 h rt, washed with 20 column volumes PBS + 20 mM imidazole, and eluted twice with 5 column volumes PBS + 250 mM imidazole. Antibodies and AbLecs were buffer exchanged into PBS using PD-10 desalting columns (GE). Purified avelumab monoclonal antibody was purchased from Selleck Chemicals.

### AbLec gel characterization

For SDS-PAGE gels, 2 μg of protein with SDS loading dye (non-reducing conditions), or SDS dye + 1 M betamercaptoethanol, heated at 95 °C for 5 min (reducing conditions), were loaded onto a Criterion™ XT 4-12% Bis-Tris Protein Gel, 18 well, (Bio-Rad) and run with XT-MES buffer at 180 V for 40 min. Total protein content was visualized using Aquastain (Bulldog Bio) with staining for 10 minutes followed by a 10 minute destain in water. For AbLec Western blots, 0.2 μg of protein was loaded onto an SDS-PAGE gel and run as described above, transferred to a nitrocellulose membrane using the Trans-Blot® Turbo™ RTA Midi Nitrocellulose Transfer Kit (Bio-Rad) at 25V, for 14 min. The membrane was blocked in Intercept® PBS blocking buffer (LI-COR) for 0.5-1 h rt, then stained with anti-HA Tag Polyclonal Antibody (clone SG77, Thermo Fisher Scientific; 1:500) and anti-6xHis antibody (clone J099B12, BioLegend; 1:2000) for 1 hour in Intercept® PBS blocking buffer with shaking at rt. The membrane was washed 3× in PBST (PBS + 0.1% Tween), followed by staining with secondary antibodies IRDye® 800CW Goat anti-Mouse and IRDye® 680RD Goat anti-Rabbit (LI-COR; 1:15,000) in PBST for 0.5-1 h with shaking at room temp, followed by 3× washes in PBST before imaging. All gels and blots were imaged on an Odyssey® CLx Imaging System (LI-COR).

### AbLec mass spectrometry characterization

Four micrograms of each AbLec were digested with trypsin for proteomic analysis. Samples were incubated for 15 minutes at 55 °C with 5 mM dithiothreitol, followed by a 30 minute incubation at room temperature in the dark with 15 mM iodoacetamide. Trypsin (Promega) was added at a 1:20 w:w ratio and digestions proceeded at room temperature overnight. The following morning, samples were desalted by quenching the digestion with formic acid to a final pH of ∼2, followed by desalting over a polystyrene-divinylbenzene solid phase extraction (PS-DVB SPE) cartridge (Phenomenex). Samples were dried with vacuum centrifugation following desalting and were resuspended in 0.2% formic acid in water at 0.5 µg per µL.

Approximately 1 µg of peptide was injected per analysis, wherein peptides were separated over a 25 cm EasySpray reversed phase LC column (75 µm inner diameter packed with 2 μm, 100 Å, PepMap C18 particles, Thermo Fisher Scientific). The mobile phases (A: water with 0.2% formic acid and B: acetonitrile with 0.2% formic acid) were driven and controlled by a Dionex Ultimate 3000 RPLC nano system (Thermo Fisher Scientific). Gradient elution was performed at 300 nL/min. Mobile phase B was held at 0% over 6 min, followed by an increase to 5% at 7 minutes, 25% at 66 min, a ramp to 90% B at 70 min, and a wash at 90% B for 5 min. Flow was then ramped back to 0% B at 75.1 minutes, and the column was re-equilibrated at 0% B for 15 min, for a total analysis time of 90 minutes. Eluted peptides were analyzed on an Orbitrap Fusion Tribrid MS system (Thermo Fisher Scientific). Precursors were ionized using an EASY-Spray ionization source (Thermo Fisher Scientific) source held at +2.2 kV compared to ground, and the column was held at 40 °C. The inlet capillary temperature was held at 275 °C. Survey scans of peptide precursors were collected in the Orbitrap from 350-1350 m/z with an AGC target of 1,000,000, a maximum injection time of 50 ms, and a resolution of 60,000 at 200 m/z. Monoisotopic precursor selection was enabled for peptide isotopic distributions, precursors of z = 2-5 were selected for data-dependent MS/MS scans for 2 seconds of cycle time, and dynamic exclusion was set to 30 seconds with a ±10 ppm window set around the precursor monoisotope. An isolation window of 1 m/z was used to select precursor ions with the quadrupole. MS/MS scans were collected using HCD at 30 normalized collision energy (nce) with an AGC target of 100,000 and a maximum injection time of 54 ms. Mass analysis was performed in the Orbitrap a resolution of 30,000 at 200 m/z and scan range set to auto calculation.

Raw data were processed using Byonic^81^ in MaxQuant version 3.11.3. Oxidation of methionine (+15.994915) was set as a common^82^ variable modification, protein N-terminal acetylation (+42.010565) and asparagine deamination (+0.984016) were specified as rare variable modifications, and carbamidomethylation of cysteine (+57.021464) was set as a fixed modification. Up to three common and two rare modifications were permitted. A precursor ion search tolerance of 10 ppm and a product ion mass tolerance of 20 ppm were used for searches, and three missed cleavages were allowed for full trypsin specificity. Peptide spectral matches (PSMs) were made against custom FASTA sequence files that contained appropriate combinations of Siglec-7 and -9 holes, and trastuzumab knobs and light chains. Peptides were filtered to a 1% false discovery rate (FDR) and a 1% protein FDR was applied according to the target-decoy method. All peptide identifications were manually inspected, and sequences coverages were calculated only from validated peptide identifications. Sequence coverage percentages are derived from the proportion of amino acids explained by peptide identifications relative to the total number of amino acids.

### Lectin flow cytometry

Cells were pelleted by centrifugation, washed once with phosphate-buffered saline (PBS), and resuspended in FACS buffer (PBS with 0.5% bovine serum albumin). Siglec-Fc or human IgG1 isotype Fc (R&D Systems) precomplexes with Alexa Fluor 647 (AF647)-labeled donkey anti-human IgG antibody (Jackson ImmunoResearch) were prepared at equimolar concentrations (7.5 nM) in FACS buffer. Biotin-labeled MALII lectin (Vector Labs, 10 μg/mL) was precomplexed with AF647-Streptavidin (Thermo Fisher Scientific, 1 μg/mL) in FACS buffer. Staining was performed for 30 min on ice at 1 × 10^6^ cells/mL. Cells were subsequently washed twice with ice-cold FACS buffer, resuspended in FACS buffer containing 100 nM SytoxGreen or 1 μM SytoxBlue (Thermo Fisher Scientific), and analyzed by flow cytometry (BD LSR II).

### Antibody flow cytometry

Cells were pelleted by centrifugation, washed once with phosphate-buffered saline (PBS), and resuspended in FACS buffer, as above. To quantify HER2, CD20, or EGFR expression, cells were stained with 7.5 nM antibody (trastuzumab, rituximab, or cetuximab, respectively) or a human IgG1 isotype control in FACS buffer for 30 min on ice at 1 × 10^6^ cells/mL. Cells were subsequently washed twice with ice-cold FACS buffer, resuspended in FACS buffer with 7.5 nM AF647-labeled donkey anti-human IgG secondary antibody (Jackson ImmunoResearch) and stained for 30 min on ice. Following incubation with the secondary antibody, cells were washed twice with ice-cold FACS buffer, resuspended in FACS buffer containing 100 nM SytoxGreen or 1 μM SytoxBlue (Thermo Fisher Scientific), and analyzed via flow cytometry (BD LSR II). For all other flow experiments, cells were stained with 10 μg/mL dye-labeled antibodies or the appropriate isotype controls in FACS buffer for 30 min at 4 °C. All antibody clones used are listed in **Supplemental Table 2**. Following incubation, cells were washed twice with ice-cold FACS buffer, resuspended in FACS buffer containing 100 nM SytoxGreen or 1 μM SytoxBlue (Thermo Fisher Scientific), and analyzed via flow cytometry (BD LSR II).

### Quantifying AbLec binding and K_D_

HER2+ K562 and SK-BR-3 cells were isolated from the cell culture supernatant or via dissociation with TrypLE (Gibco), respectively, washed with 1xPBS, and resuspended in blocking buffer. 60,000 cells were then distributed into wells of a 96-well V-bottom plate (Corning). Various concentrations of trastuzumab, nivolumab, pembrolizumab, Siglec-Fcs (R&D Systems), Galectin-9-Fc (Sino Biological), or AbLecs were added to the cells in equal volumes and incubated with cells for 1 h at 4 °C with periodic pipet mixing. Cells were washed three times in FACS buffer (PBS + 0.5% bovine serum albumin), pelleting by centrifugation at 300 xg for 5 min at 4 °C between washes. Cells were resuspended in 4 μg/mL AF647-labeled donkey anti-human secondary antibody (Jackson ImmunoResearch) in FACS buffer for 30 min at 4 °C. Cells were further washed twice and resuspended in FACS buffer, and fluorescence was analyzed by flow cytometry (BD LSR II). Gating was performed using FlowJo v.10.0 software (Tree Star) to eliminate debris and isolate single cells. Mean fluorescence intensity (MFI) of the cell populations were normalized to the MFI of cells stained with 50 nM trastuzumab from each experimental replicate. MFI values were fit to a one-site total binding curve using GraphPad Prism 9.0, which calculated the *K*_D_ values as the antibody concentration needed to achieve a half-maximum binding.

### Computational modeling of AbLec binding and apparent ligand concentrations

The *MVsim* multivalent simulation application (version 1.1 with MATLAB 2021a) was used to model trastuzumab, T7 AbLec, and T7A AbLec binding to HER2+ K562 cells, as previously reported^42^. *MVsim* models the effects of molecular valency, topology, affinity, and kinetics on the binding dynamics of multivalent and multispecific macromolecular receptor-ligand systems. Here, the *MVsim* application was used to create a modeling routine that simulated, via the microstate binding model^42^, the interaction of antibodies and antibody-like chimeras to surfaces populated with specific targeted ligands/antigens. The designed sequences and structures of the biomolecules were used to parameterize the model with the binding valencies. Here, the trastuzumab, T7A AbLec, and Siglec-7-Fc reference biomolecules, as well as the T7 AbLec, were modeled as topologically related, Fc domain-driven, immunoglobulin dimers, each with a valency of two. The model was further parameterized with monovalent binding affinities (*K*_D_ values) that were experimentally determined here (as described in the above Methods; *Quantifying AbLec binding and K_D_*) and informed by the supporting literature. Affinities were parameterized as follows: trastuzumab monovalent *K*_D_ = 11.2 nM and Siglec-7-Fc monovalent *K*_D_ = 200 µM^83,84^.

The parameterized model was then used to fit the flow cytometry-based experimental binding data for the apparent concentration ([C_app_]) parameters that derive from the spatial, nearest-neighbor proximity of pairs of targeted cell surface ligands/antigens. Model-derived estimates of [C_app_] for various designed topologies were interpreted as quantitative assessments of the relative probability or favorability of a candidate therapeutic engaging with a particular cell surface in an advantageous bivalent and high-avidity interaction.

### Expression and dye labeling of Siglec-Fc reagents

Siglec-Fc decoy receptors were expressed in Expi293F cells that stably express human formylglycine-generating enzyme for aldehyde tagging. Siglec-Fc constructs were expressed according to the Expi293F manufacturer protocol (Thermo Fisher Scientific) and purified by manual gravity column (BioRad) using protein A agarose (Fisher Scientific), as described for antibodies above, with the modification that transfected cells, supernatant, and purified Siglec-Fcs were stored in the dark. Siglecs were then buffer exchanged into acidic buffer, concentrated, and conjugated with HIPS-azide as we previously described^16^. Siglec-Fc-azide was then buffer exchanged into PBS, and 100× molar equivalents of DBCO-AF647 (Click Chemistry Tools, 1302-1) in DMSO were added and the reaction was mixed at 500 rpm in the dark for 2 hours rt. Siglecs were buffer exchanged into PBS by 6× centrifugation on Amicon columns (30 kDa MWCO) and AF647 addition was confirmed by using a NanoDrop spectrophotometer at 650 nm for the AF647 dye (extinction coefficient 239,000) and at 280 nm for the protein (using extinction coefficients calculated for each Siglec by Expasy).^85^

### Competitive binding assays

HER2+ K562 cells were isolated from the cell culture supernatant, washed with 1xPBS, and resuspended in blocking buffer. Cells were aliquoted for sialidase treatment with 100 nM *V. cholerae* sialidase at 37 °C for 30 min in blocking buffer. 60,000 untreated or sialidase treated cells were then distributed into wells of a 96-well V-bottom plate (Corning). Various concentrations of trastuzumab or AbLecs were combined with 200 nM Siglec-7/9-Fc-AF647 or 0.3 μg/mL MEM-59-AF647, added to the cells in equal volumes and incubated with cells for 2-3 h at 4 °C with periodic pipet mixing. Cells were washed three times and resuspended in blocking buffer, pelleting by centrifugation at 300 xg for 5 min at 4 °C between washes, and fluorescence was analyzed by flow cytometry (BD LSR II). Gating was performed using FlowJo v.10.0 software (Tree Star) to eliminate debris and isolate single cells. Mean fluorescence intensity (MFI) of the cell populations were background subtracted and normalized to the MFI of cells stained with 200 nM Siglec-7/9-Fc-AF647 without antibody or AbLecs from each experimental replicate.

### Cell growth and toxicity assays

SK-BR-3 and HER2+ K562 cells were lifted with 2 mL trypsin for 5 min at 37 °C, rinsed with 8 mL normal growth media, pelleted by centrifugation at 300 *x g*, and resuspended in phenol-red-free growth medium containing 50 nM Sytox green cell dead stain (Thermo Fisher Scientific) for HER2+ K562 cells or 5 nM Sytox red dead cell stain (Thermo Fisher Scientific) for SK-BR-3 cells to measure cytotoxicity. Cells were plated onto a flat-bottomed 96 well plate (10,000 cells per well, 95 μL), then 5 μL of AbLec, Siglec, or antibody in PBS was added and mixed, followed by centrifugation at 30 *x g* for 1 min. Images were acquired using an IncuCyte® S3 Live-Cell Analysis System (Sartorius) every 2 h for 3 days. SK-BR-3 cells were analyzed by phase with segmentation adjustment = 1, and a minimum area of 200 μm^2^, cell death was quantified by red fluorescence using a threshold of 0.3 RCU, with an edge sensitivity of -50 and areas between 50 and 1000 square microns. HER2+ K562 phase segmentation adjustment was 0.2, with no hole-fill and a minimum area of 60 microns. Cell death was quantified by green fluorescence using a threshold of 2 RCU, with an edge sensitivity of -45 and areas between 50 and 800 microns with eccentricity and integrated intensities less than 0.95 and 40000, respectively.

### Isolation and differentiation of donor macrophages

LRS chambers were obtained from anonymous healthy donors through the Stanford Blood Center. PBMCs were isolated using Ficoll-Paque (Sigma-Aldrich) density gradient centrifugation. Isolated PBMCs were extracted from the PBS/Ficoll interface, washed once with PBS, and resuspended in serum-free RPMI. Monocytes were isolated by plating ∼1×10^8^ PBMCs in a T75 flask containing serum-free RPMI for 1-2 hours, followed by 3× washes with PBS +Ca +Mg to remove non-adherent cells. The media was then replaced with IMDM with 10% Human AB Serum (Gemini) and incubated for 7-9 days to induce macrophage differentiation prior to use in phagocytosis or flow cytometry experiments.

### Phagocytosis assays

Macrophages were washed with PBS and lifted by 30 min incubation at 37 °C with 10 mL TrypLE (Thermo Fisher Scientific). Macrophages were pelleted by centrifugation at 300 *x g* for 5 min and resuspended in IncuCyte medium (phenol-red free RPMI + 10% HI FBS). Macrophages (10,000 cells, 100 μL) were added to a 96 well flat-bottom plate (Corning) and incubated in a humidified incubator for 1 h at 37 °C. Meanwhile, target cells were washed 1× with PBS, then treated with 1:80,000 diluted pHrodo red succinimydyl ester dye (Thermo Fisher Scientific) in PBS at 37 °C for 30 min, washed 1× with PBS and resuspended in IncuCyte medium. Finally, 10 μL of 20× antibody or AbLec stocks in PBS were added to the macrophages, followed by addition of pHrodo red-stained target cells (20,000 cells, 90 μL). Antibodies, Siglec-Fcs, and AbLecs were added at 25 nM, Siglec-7 and -9 blocking antibodies (1E8 and mAbA clones, respectively^20^; Creative Biolabs) were used at 12.5-25 μg/mL, *V. cholerae* sialidase was used at 100 nM, and human IgG1 isotype antibody (BioXCell), rat IgG2a isotype antibody (BioXCell), and CD47 blocking antibody (clone B6H12, BioXCell)^23^ was added at 10 μg/mL unless otherwise noted. Cells were plated by gentle centrifugation (50 *x g*, 2 min). Two images per well were acquired using an IncuCyte® S3 Live-Cell Analysis System (Sartorius) at 1 h intervals until the maximum signal was reached. The quantification of pHrodo red fluorescence was empirically optimized for phagocytosis of each cell line based on their background fluorescence and size. K562 and MDA-MB-231 cells were analyzed with a threshold of 0.8, an edge sensitivity of -70, and the area was gated to between 100 and 2000 μm^2^ with integrated intensities between 300 and 2000 RCU x μm^2^ / image. HCC-1954 were analyzed with a threshold of 1.5, an edge sensitivity of -45, and an area between 30 and 2000 μm^2^. SK-BR-3 analysis had a threshold of 1, an edge sensitivity of -55, a minimum integrated intensity of 60, and a maximum area and eccentricity 3000 and 0.96, respectively. Ramos and Raji analysis used a threshold of 1.5, an edge sensitivity of -45, and areas between 100 and 2000 μm^2^. The total red object integrated intensity (RCU × µm²/Image) was quantified for each image and the maximum total red object integrated intensity value is reported for each treatment condition.

### Isolation of donor NK cells

PBMCs were isolated from LRS chambers as described above, cryostocks were prepared at 2-4×10^7^ cells in 90% heat-inactivated FBS + 10% DMSO and stored in liquid nitrogen vapor until use. The day prior to use stocks were thawed, NK cells were isolated using an EasySep NK isolation kit (StemCell Technologies), and cells were cultured with 0.5 µg/mL recombinant IL-2 (BioLegend) in RPMI + 10% heat-inactivated FBS for 24 h before use in NK cell cytotoxicity or flow cytometry experiments.

### NK cell killing assays

Target cells were lifted with TrypLE (Thermo Fisher Scientific) and stained with CellTracker Deep Red dye (Thermo Fisher Scientific) according to the manufacturer’s protocol. NK cells and target cells were mixed at an effector to target (E/T) ratio of 4:1 (SK-BR-3 cells) or 2:1 (K562 cells) and Sytox Green (Thermo Fisher Scientific) was added at 100 nM. Antibodies, Siglec decoy receptors, and AbLecs were added at 25 nM, Siglec-7 and -9 blocking antibodies (1E8 and mAbA clones, respectively; Creative BioLabs)^20^ were used at 12.5-25 μg/mL, CD16 blocking antibody (clone 3G8, BioLegend) was added at 10 μg/mL, and *V. cholerae* sialidase was used at 100 nM, unless otherwise noted. After 5-6 h incubation at 37 °C, cell death was analyzed by flow cytometry by selecting the CellTracker Deep Red^+^ cells and quantifying the percent dead as Sytox Green^+^ / total CellTracker Deep Red^+^ cells. NK cytotoxicity index was then calculated as the ratio of dead to alive cells in the assay: % dead / (100-% dead).

### Isolation of human PMNs

Polymorphonuclear granulocytes (PMN) were isolated from peripheral blood of healthy donors by density gradient centrifugation using either Polymorphprep^®^ (Progen, Heidelberg, DE), as previously described^86^. Isolated PMNs were >82% pure by flow cytometry.

### PMN cytotoxicity assays

PMN-mediated cytotoxicity was analyzed in chromium-51 [^51^Cr] release assays as previously described^87^. AbLecs and antibodies were added at varying concentrations. The CD47 blocking antibody hu5F9-IgG2σ was applied at 20 µg/ml. Effector cells and ^51^Cr-labelled target cells were added at a ratio of 40:1. After 3 h incubation at 37 °C, ^51^Cr release was measured in counts per minute (cpm) in a MikroBetaTrilux 1450 liquid scintillation and luminescence counter (PerkinElmer, Rodgau Jügensheim, DE). Maximal ^51^Cr release was achieved by addition of 2% v/v Triton-X 100 solution, while basal ^51^Cr release was measured in the absence of antibodies. Specific tumor cell lysis in % was calculated as follows:

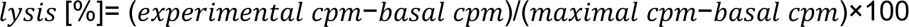

### Acquisition of IncuCyte images

For phagocytosis and cytotoxicity assays, images were obtained over time using an Incucyte® S3 Live-Cell Analysis System (Essen BioScience) within a Thermo Fisher Scientific tissue culture incubator at maintained 37 °C with 5% CO_2_. Data were acquired from a 10× objective lens in phase contrast, a green fluorescence channel (ex: 460 ± 20, em: 524 ± 20, acquisition time: 300 ms), and from a red fluorescence channel (ex. 585 ± 20, em: 665 ± 40, acquisition time: 400 ms). Two images per well were acquired at intervals. Unless otherwise specified, all cells were analyzed by Top-Hat segmentation with 100 μm radius, edge split on, hole fill: 0 μm^2^.

### Mice

WT C57BL/6 mice were purchased from The Jackson Laboratories and housed at Stanford University. Experiments involving animals at Stanford University were approved by the Administrative Panel on Lab Animal Care (APLAC) and carried out according to protocol number 31511. Stanford University vivaria are AAALAC-accredited. Husbandry is performed in accordance with the Guide for the Care and Use of Laboratory Animals and the Public Health Service Policy on Humane Care and Use of Laboratory Animals. SigE^-/-^ and SigE^-/-^ Sig7/9 mice were developed previously^20^ and were bred and maintained at the Comparative Bioscience Center at The Rockefeller University. All experiments carried out at The Rockefeller University were performed in compliance with federal laws and institutional guidelines and have been approved by The Rockefeller University Institutional Animal Care and Use Committee. Animals were housed under the following conditions: 12-h light/12-h dark cycle, ambient temperature of 20–22 °C and relative humidity of between 30 and 70%. All studies were performed on 8- to 12-wk old age- and sex-matched female and male mice.

### Isolation of murine bone marrow

Femurs, tibias, and fibias were collected from WT, SigE^-/-^, and SigE^-/-^ Sig7/9 mice. Bones were sterilized with 96% ethanol and rinsed with PBS. Proximal and distal ends of bones were removed and a 1 mL syringe with 26 gauge needle containing PBS was used to extract bone marrow and resuspend until an approximately single cell suspension was achieved. The cell suspension was then passed over a 70 µm filter, cells were pelleted at 200-250 xg for 5-10 min at rt, resuspended at 5 x 10^6^ cells/mL in heat-inactivated FBS + 10% DMSO, and stored in liquid nitrogen vapor until use.

### Murine BMDM differentiation and activation assays

Murine bone marrow cryostocks were thawed into 10 mL of warmed complete media (RPMI + 10% HI FBS + 1% penicillin/streptomycin), pelleted at 300 xg for 5 min rt, and plated in complete media + 0.1 µg/mL M-CSF (PeproTech). After 3 and 5 days of incubation, the media was replaced with complete media + 0.1 µg/mL M-CSF (PeproTech). BMDMs were allowed to differentiate in refreshed media until use on days 7-9. For use in activation assays, BMDMs were lifted by 20 min incubation at 37 °C with 10 mL TrypLE (Thermo Fisher Scientific). BMDMs (10-15,000 cells, 100 μL) in complete media + 0.1 µg/mL M-CSF were added to a 96 well flat-bottom plate (Corning) and incubated in a humidified incubator for 1 h at 37 °C. Meanwhile, HER2+ B16D5 cells were washed 1× with PBS, treated with 1:80,000 diluted pHrodo red succinimydyl ester dye (Thermo Fisher Scientific) in PBS at 37 °C for 30 min, washed 1× with PBS and resuspended in complete media + 0.1 µg/mL M-CSF. Finally, 10 μL of 20× antibody or AbLec stocks in PBS were added to the macrophages, followed by addition of pHrodo red-stained target cells (20,000 cells, 90 μL) and BMDM/target cell co-cultures were incubated for 24 h. Following incubation, 100 μL supernatant was collected and analyzed using a Cytometric Bead Array Mouse Inflammation Kit (BD), according to the manufacturer’s protocol.

### Isolation and activation of human T cells

PBMCs were isolated from LRS chambers, cryostocks were prepared at 2-4×10^7^ cells in 90% heat-inactivated FBS + 10% DMSO, and stocks were stored in liquid nitrogen vapor until use, as described above. Human T cells were isolated from thawed cryostocks using an EasySep CD3+ T cell isolation kit (StemCell Technologies). T cells were cultured at a 1:2 cell:bead ratio with cultured αCD3/αCD28 activator beads (Thermo Fisher Scientific) and 50 ng/mL recombinant IL-2 (BioLegend) in RPMI + 10% heat-inactivated FBS for 3 days before use.

### Statistical Analysis

Statistical analysis was performed in Prism (version 9.0). For binding curves, one-site-specific binding curves were used to calculate the dissociation constants of antibodies and AbLecs. In binding assays, NK cell cytotoxicity experiments, phagocytosis experiments, and Siglec expression analyses, ordinary one-way ANOVAs were performed with Tukey’s multiple comparison’s test to compare treatment groups. All results were confirmed with multiple biological replicates as indicated in the figure legends. All data were tested for outliers using GraphPad Outlier Calculator (http://graphpad.com/quickcalcs/Grubbs1.cfm) and no outliers were identified. In every instance ns indicates p>0.05, * indicates p<0.05, ** indicates p<0.01, *** indicates p<0.001, and **** indicates p<0.0001.

## Acknowledgements

J.C.S. acknowledges support from NIH/NCI F32 Postdoctoral Fellowship 1F32CA250324-01, American Cancer Society Postdoctoral Fellowship PF-20-143-01-LIB, and a Sarafan ChEM-H Postdocs at the Interface seed grant. S.W. is supported by funding from the Canadian Institutes of Health Research, the Natural Sciences and Engineering Research Council of Canada, the Cancer Research Society, the Canadian Cancer Society, the Arthritis Society of Canada, Health Research BC, and the Canadian Glycomics Network (GlycoNet). M.L. and T.V. acknowledge funding from the German Research Foundation (DFG KFO 5010 P6). M.A.G. acknowledges support from the National Science Foundation Graduate Research Fellowship and the Sarafan ChEM-H Chemistry/Biology Interface Predoctoral Training Program. N.M.R. acknowledges support from NIH grant K99GM147304. C.A.S. acknowledges support from NIH (R35 GM136309). C.R.B. acknowledges support from NIH (R01 GM058867-23, R01 CA227942, and U01 CA226051-02) and a Merck Research Labs Discovery Biologics SEEDS grant. Cell sorting/flow cytometry analysis for this project was performed using instruments in the Stanford Shared FACS Facility.

## Author Contributions

J.C.S., M.A.G., S.P.W., and C.R.B. conceived the study. J.C.S., M.A.G., S.P.W., I.I.B., N.M.R., M.K.R, M.L., W.J.E., and B.B. designed and performed experiments and analyzed data. C.A.S., J.V.R., T.V., and C.R.B. directed research and aided in data analysis. J.C.S., M.A.G., and C.R.B. wrote the manuscript. All authors reviewed and/or revised the manuscript.

## Data and Materials Availability Statement

All primary data for main text and supplementary figures are available from the authors upon reasonable request. Any unique materials presented in the manuscript may be available from the authors upon reasonable request and through a materials transfer agreement.

