## Supplemental Information for "Antibody-lectin chimeras for glyco-immune checkpoint blockade"

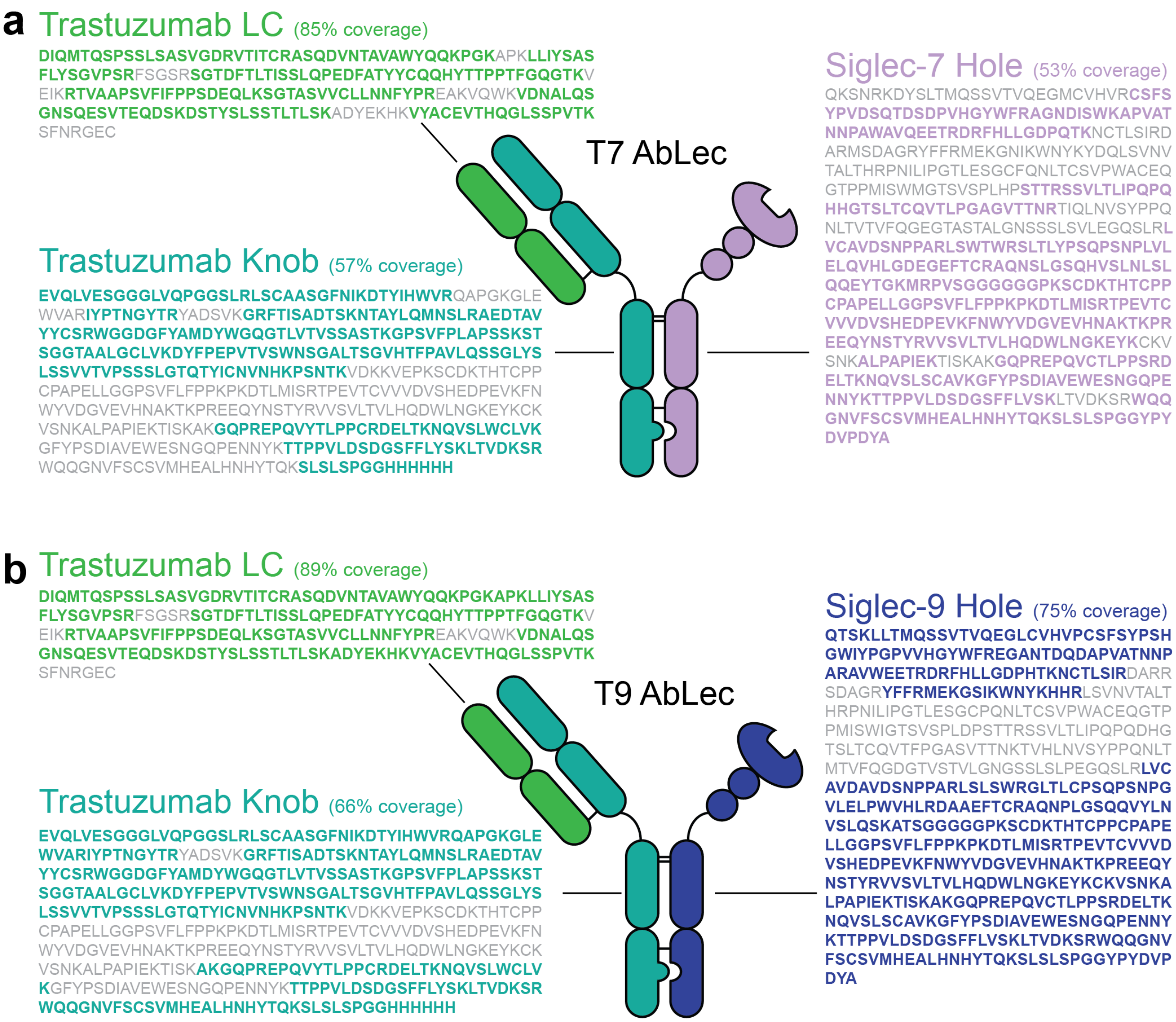


**Extended Data Figure 1. Mass spectrometry sequencing of T7 and T9 AbLecs.**

Proteomic analysis of (**a**) T7 and (**b**) T9 AbLecs provides evidence of expression of trastuzumab heavy chain, light chain, and Siglec-Fc decoy receptor chain with similar levels of sequence coverage.

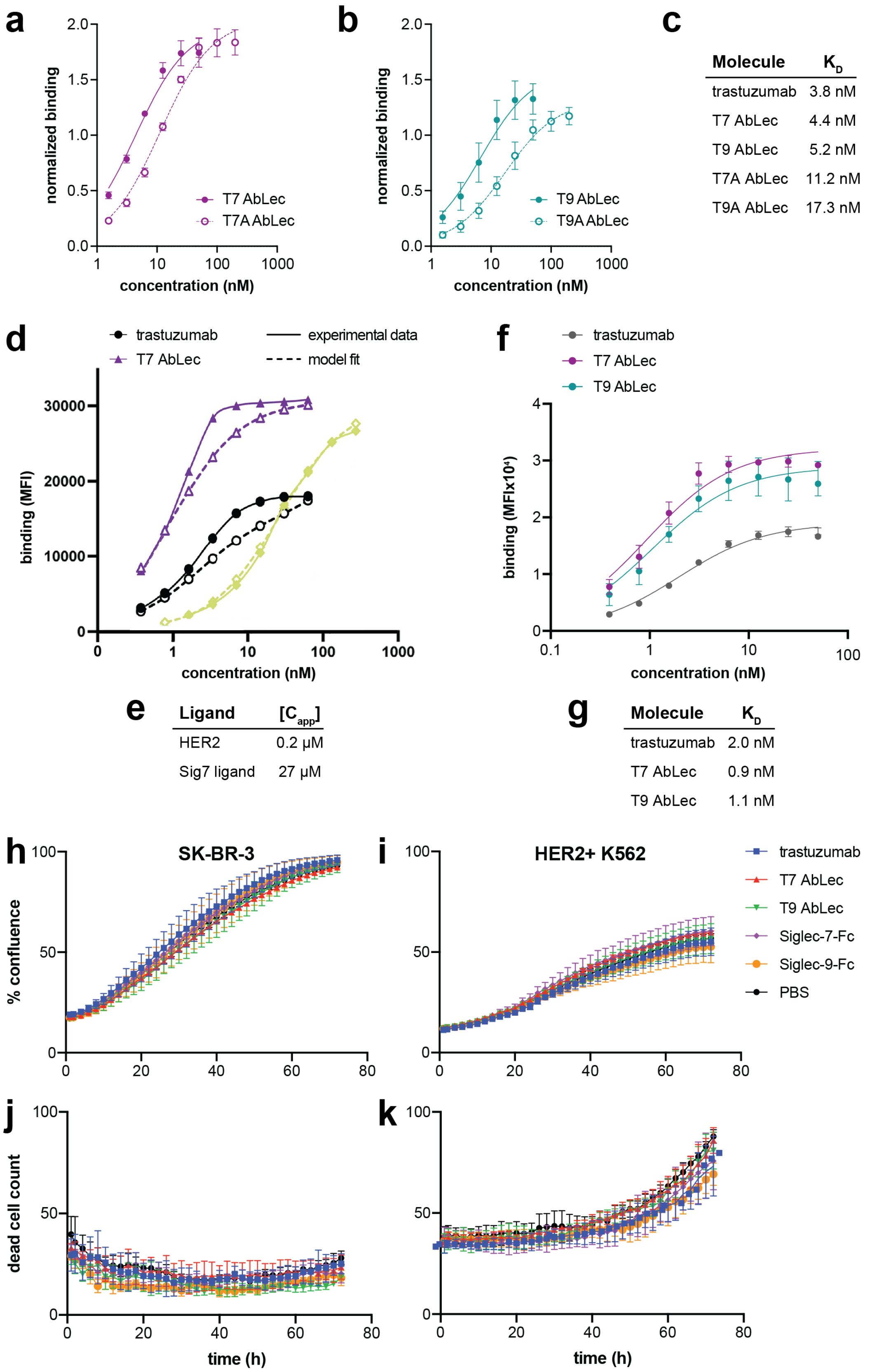
a
normalized binding
concentration (nM)
T7 AbLec
T7A AbLec
b
normalized binding
concentration (nM)
T9 AbLec
T9A AbLec
c
Molecule | $K_D$
trastuzumab | 3.8 nM
T7 AbLec | 4.4 nM
T9 AbLec | 5.2 nM
T7A AbLec | 11.2 nM
T9A AbLec | 17.3 nM
d
trastuzumab
T7 AbLec
experimental data
model fit
binding (MFI)
concentration (nM)
e
Ligand | $[C_{app}]$
HER2 | 0.2 μM
Sig7 ligand | 27 μM
f
trastuzumab
T7 AbLec
T9 AbLec
binding (MFIx10⁴)
concentration (nM)
g
Molecule | $K_D$
trastuzumab | 2.0 nM
T7 AbLec | 0.9 nM
T9 AbLec | 1.1 nM
h
SK-BR-3
% confluence
i
HER2+ K562
trastuzumab
T7 AbLec
T9 AbLec
Siglec-7-Fc
Siglec-9-Fc
PBS
j
dead cell count
time (h)
k
time (h)

**Extended Data Figure 2. AbLec binding displays higher avidity and cooperativity than trastuzumab due to high ratio of Siglec ligand expression to targeted antigen expression.**

(**a, b**) Binding of T7 and T9 AbLecs, or R→A AbLec mutants with reduced Siglec binding (T7A and T9A AbLecs) to SK-BR-3 cells was quantified via flow cytometry (n = 3, binding normalized to 50 nM trastuzumab for each experimental replicate).

(**c**) Dissociation constants for trastuzumab, T7, T9, T7A, and T9A AbLec binding determined by fitting experimental data from (**a**) to a one-site total binding curve.

(**d**) Binding of trastuzumab, T7 AbLec, and T7A AbLec to HER2+ K562 cells was modeled using *MVsim*[42]. A modeling routine was constructed to simulate the enhancement to binding affinity ($K_D$) that was achieved through the cooperative binding of either a bivalent, monospecific antibody (trastuzumab; black traces) or a bivalent, bispecific AbLec (T7 AbLec; purple traces) to a surface consisting of both HER2 and Siglec-7 ligands. Cooperative binding was assessed relative to the binding of a non-cooperative, monovalent reference (T7A AbLec; green traces).

(**e**) The *MVsim* model of T7 AbLec binding was used to estimate the apparent concentrations ([$C_{app}$]) that drive cooperative ligand binding to HER2 antigens and Siglec-7 ligands on HER2+ K562 cells. The modeling routine in (**d**) treated the avidity enhancements relative to the monovalent reference as fitted parameters from which the spatial proximities of either two HER2 antigens or one HER2 antigen and one Siglec-7 ligand at the HER2+ K562 cell surface could be estimated. These estimates are presented as [$C_{app}$] concentrations, whereby a higher [$C_{app}$] indicates a higher probability that an antibody or AbLec can engage in a cooperative, bivalent interaction at the cell surface due to proximity of its two antigens/ligands.

(**f**) Binding of trastuzumab, T7, and T9 AbLecs (T7A and T9A AbLecs) to HER2+ K562 cells was quantified via flow cytometry. Data are mean ± s.d. of n = 3 biological replicates.

(**g**) Dissociation constants for trastuzumab, T7, and T9 AbLec binding to HER2+ determined by fitting experimental data from (**f**) to a one-site total binding curve.

(**h-k**) No statistically significant difference in cell growth (**h, i**) or cell death (**j, k**) was observed in SK-BR-3 or HER2+ K562 cultures treated with T7 or T9 AbLecs compared to vehicle, trastuzumab, or Siglec decoy receptor controls. Data are mean ± s.d. of n = 3 biological replicates.

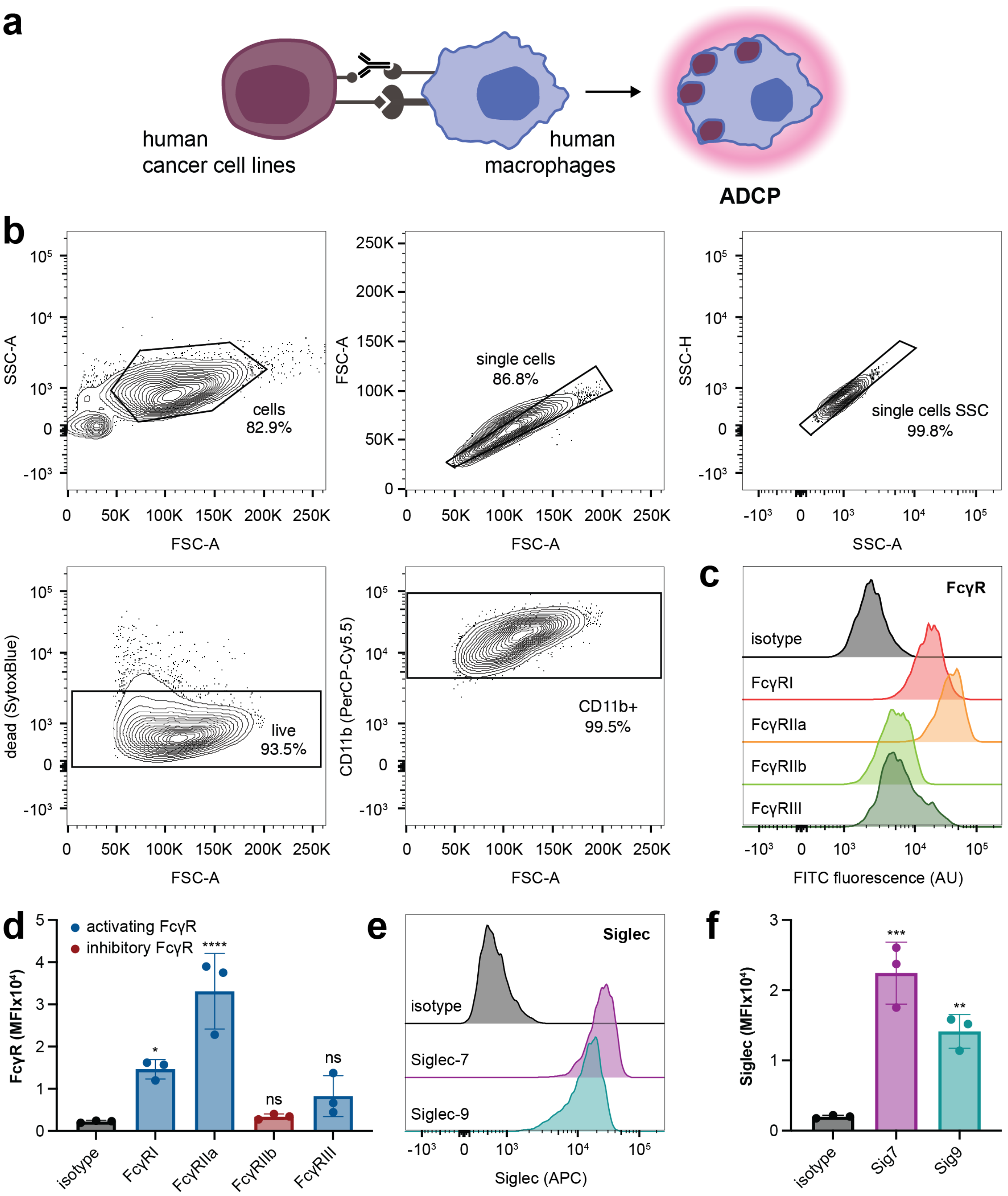


**Extended Data Figure 3. Characterization of primary human macrophages used for *in vitro* phagocytosis assays.**

(**a**) Macrophages were co-cultured with human tumor cell lines labeled with the pH-sensitive pHrodo red dye that fluoresces red in acidic phagosomes. This enabled quantification of phagocytosis via fluorescence microscopy.

(**b**) Macrophages were defined as CD11b+ by flow cytometry.

(**c**) Macrophages were profiled for expression of human Fc gamma receptors FcγRI, FcγRIIa, FcγRIIb, and FcγRIII by flow cytometry. Histograms are representative of macrophages from n = 3 human donors (quantified in **d**).

(**d**) Expression of FcγRI and FcγRIIa on macrophages measured via flow cytometry was significant compared to staining with an isotype antibody. Data are mean ± s.d. from n = 3 human donors.

(**e**) Macrophages expressed Siglec-7 and Siglec-9 receptors. Histograms are representative of macrophages from n = 3 human donors (quantified in **f**).

(**f**) Expression of Siglec-7 and Siglec-9 receptors on macrophages measured via flow cytometry was significant compared to staining with an isotype antibody. Data are mean ± s.d. from n = 3 human donors.

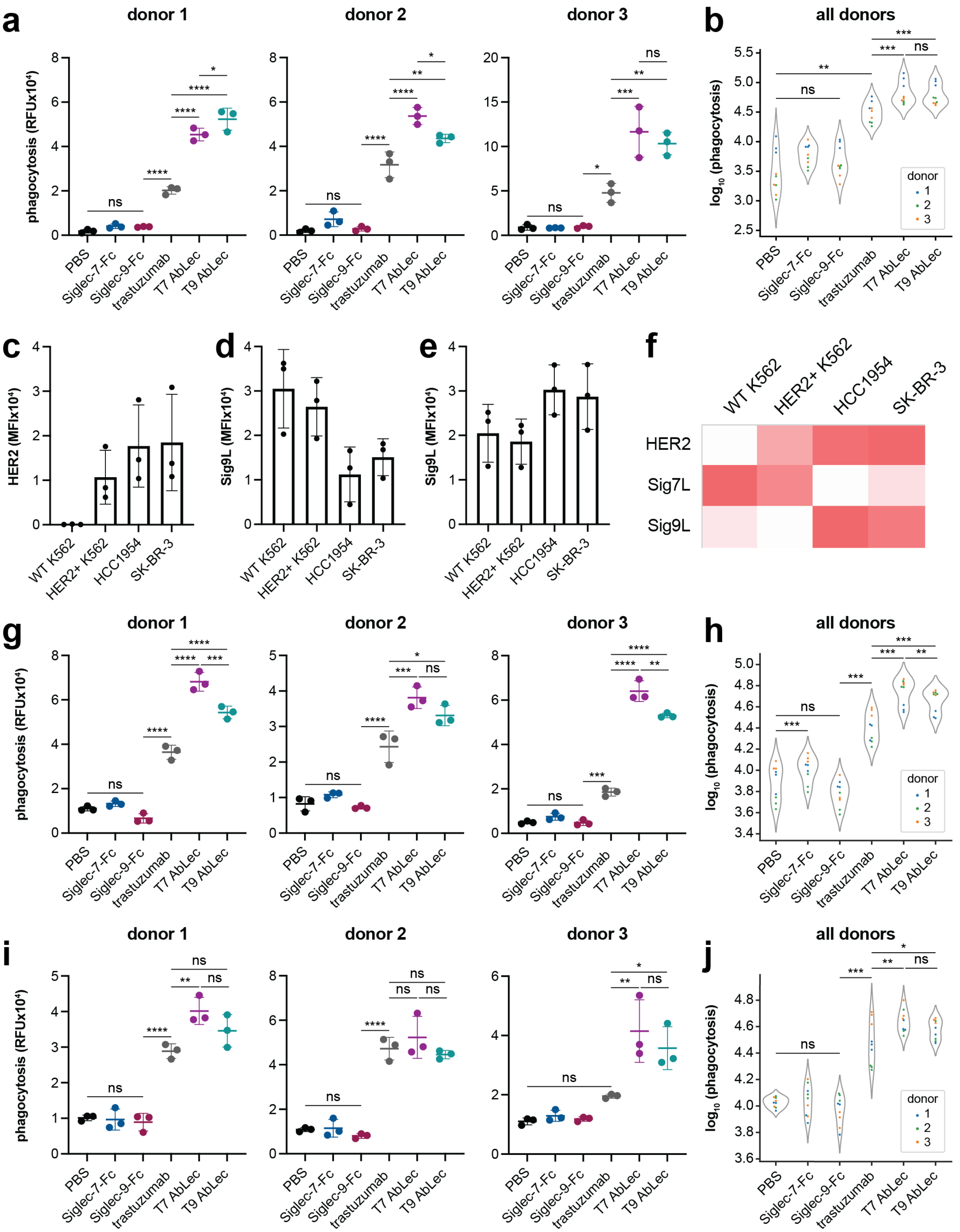


**Extended Data Figure 4. AbLec-mediated enhancement of ADCP is observed across cell lines with varying levels of HER2 and Siglec ligand expression.**

(**a**) ADCP of SK-BR-3 cells by primary human macrophages treated with T7/9 AbLecs, trastuzumab, or Siglec-7/9-Fc. Donor 3 data are also shown in **Fig. 2d**. Data are mean ± s.d. of n = 3 biological replicates.

(**b**) Statistical analysis of experimental data from (**a**) was performed using Heirarch[88] to calculate significant differences in treatment conditions across donors (n = 3 biological replicates from each of n = 3 donors).

(**c-e**) HER2+ K562, HCC-1954, and SK-BR-3 cells express varying levels and ratios of the HER2 antigen (**c**), Sig7L (**d**), and Sig9L (**e**) as measured via flow cytometry. Data are mean ± s.d. from n = 3 biological replicates.

(**f**) Heat map representation of average HER2 and Sig7/9L expression measured in (**c-e**) in HER2+ K562, HCC-1954, and SK-BR-3 cell lines compared to WT K562 cells.

(**g**) ADCP of HER2+ K562 cells by primary human macrophages treated with T7/9 AbLecs, trastuzumab, or Siglec-7/9-Fc. Data are mean ± s.d. of n = 3 biological replicates.

(**h**) Statistical analysis of experimental data from (**g**) was performed using Heirarch[88] to calculate significant differences in treatment conditions across donors (n = 3 biological replicates from each of n = 3 donors).

(**i**) ADCP of HCC-1954 cells by primary human macrophages treated with T7/9 AbLecs, trastuzumab, or Siglec-7/9-Fc. Data are mean ± s.d. of n = 3 biological replicates.

(**j**) Statistical analysis of experimental data from (**i**) was performed using Heirarch[88] to calculate significant differences in treatment conditions across donors (n = 3 biological replicates from each of n = 3 donors).

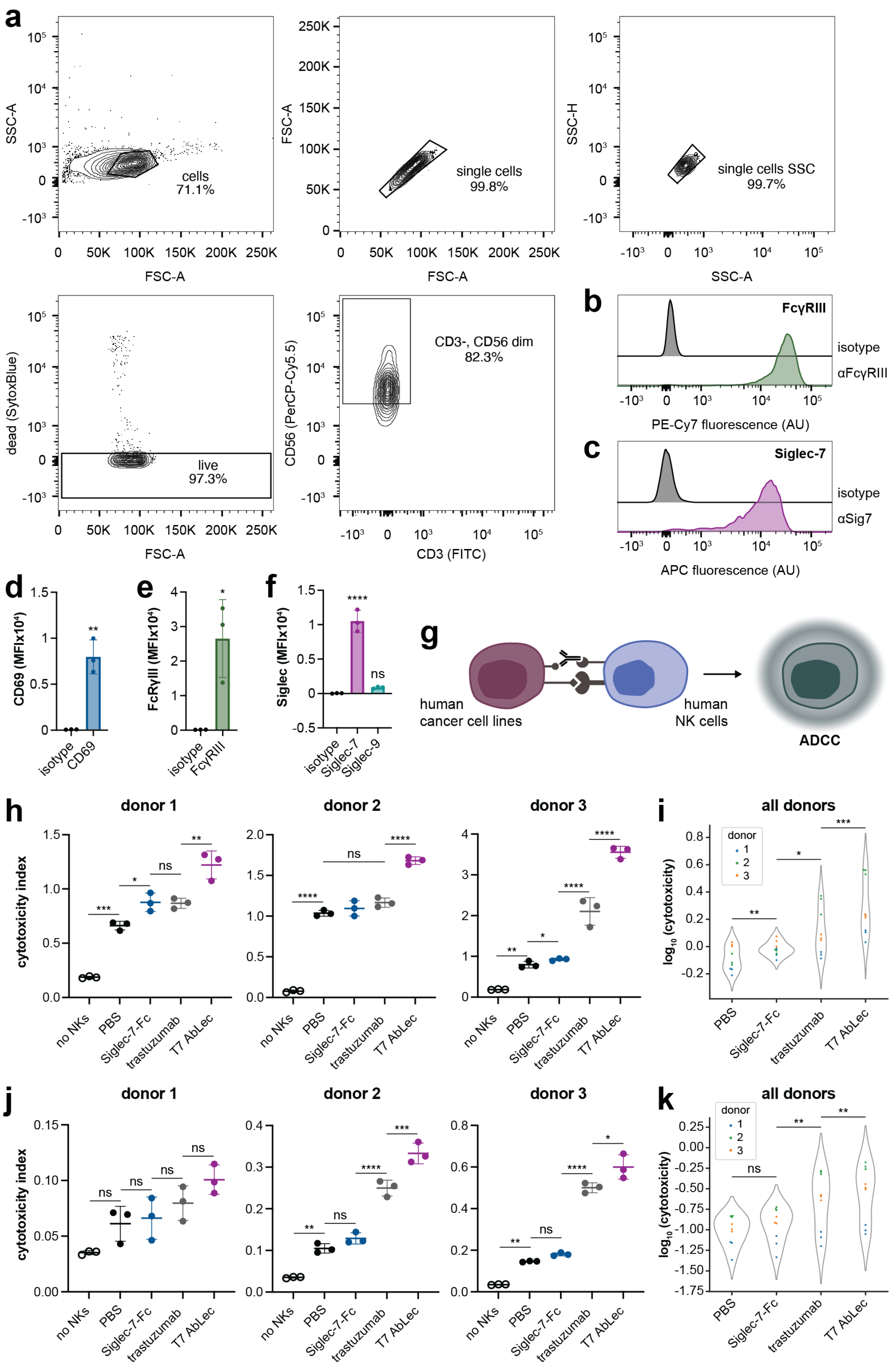


**Extended Data Figure 5. AbLecs elicit enhanced NK cell ADCC *in vitro*.**

(**a**) NK cells were defined as CD3- CD56 dim by flow cytometry.

(**b-c**) NK cells were profiled for expression of FcγRIII (**c**) and Siglec-7 (**d**) by flow cytometry. Histograms are representative of NK cells from n = 3 human donors (quantified in **e, f**).

(**d-f**) Quantification of CD69 (activation marker; **d**), FcγRIII (**e**), and Siglec-7 (**f**) expression on NK cells via flow cytometry compared to staining with an isotype antibody. Data are mean ± s.d. from n = 3 human donors (representative histograms shown in **b, c**).

(**g**) We co-cultured primary human NK cells with CellTracker Deep Red-labeled human tumor cell lines in the presence of the Sytox Green DNA intercalating dye that permeates the compromised membranes of dying cells, enabling quantification of NK cell killing of target tumor cells via flow cytometry.

(**h**) ADCC of SK-BR-3 cells by NK cells treated with T7/9 AbLecs, trastuzumab, or Siglec-7/9-Fc. Donor 3 data are also shown in **Fig. 2e**. Data are mean ± s.d. of n = 3 biological replicates.

(**i**) Statistical analysis of experimental data from (**e**) was performed using Heirarch[88] to calculate significant differences in treatment conditions across donors (n = 3 biological replicates from each of n = 3 donors).

(**j**) ADCC of HER2+ K562 cells by NK cells treated with T7/9 AbLecs, trastuzumab, or Siglec-7/9-Fc. Data are mean ± s.d. of n = 3 biological replicates.

(**k**) Statistical analysis of experimental data from (**g**) was performed using Heirarch[88] to calculate significant differences in treatment conditions across donors (n = 3 biological replicates from each of n = 3 donors).

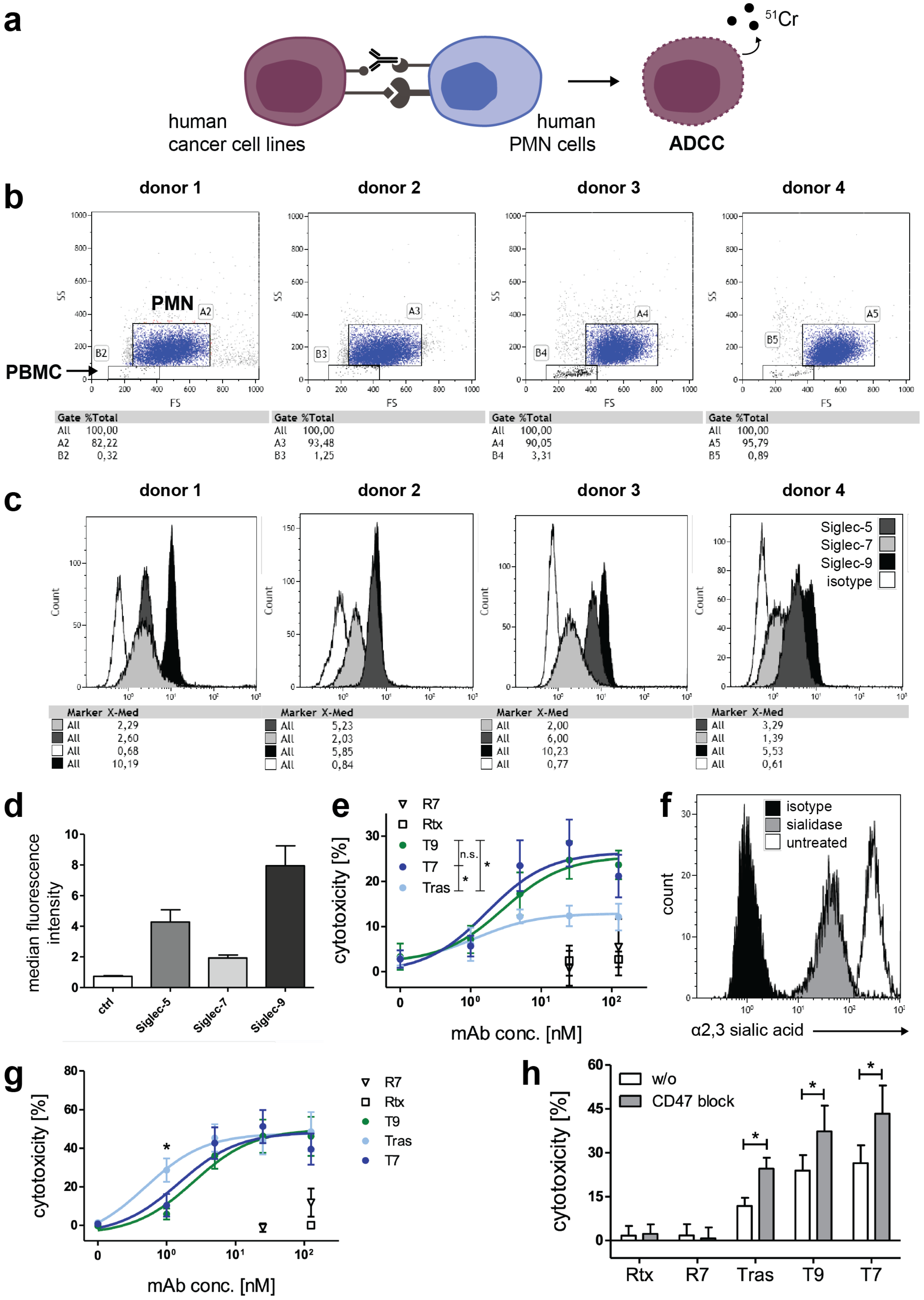

**Extended Data Figure 6. AbLecs elicit enhanced PMN ADCC *in vitro*.**

(**a**) Primary human PMNs were co-cultured with SK-BR-3 cells labeled with radioactive chromium ($^{51}Cr$). Tumor cell lysis is quantified by measuring chromium in the supernatant released by dying cells.

(**b**) PMNs were defined by distinctive forward and side scatter compared to peripheral blood mononuclear cells (PBMCs).

(**c**) PMNs from n = 4 human donors expressed the Siglec-5, -7, and -9 receptors.

(**d**) Quantification of Siglec-5, -7, and -9 expression on PMNs. Data are mean ± s.d. from n = 4 human donors.

(**e**) Quantification of PMN ADCC against HER2+ SK-BR-3 cells. Co-cultures were treated with increasing concentrations of T7/T9 AbLec, trastuzumab, isotype antibody (Rtx: rituximab), or isotype AbLec (R7: rituximab x Siglec-7) controls. Data are mean ± s.d. of replicates from n = 5 human donors.

(**f**) Cell surface α2,3 linked sialic acid expression measured via staining sialidase treated or untreated SK-BR-3 cells with the MALII lectin that detects α2,3 linked sialic acid and analysis by flow cytometry.

(**g**) Quantification of PMN ADCC against sialidase treated HER2+ SK-BR-3 cells. Co-cultures were treated with increasing concentrations of T7/T9 AbLec, trastuzumab, isotype antibody (Rtx: rituximab), or isotype AbLec (R7: rituximab x Siglec-7) controls. Data are mean ± s.d. of replicates from n = 5 human donors.

(**h**) Quantification of PMN ADCC of HER2+ SK-BR-3 cells. Co-cultures were treated with of T7/T9 AbLec, trastuzumab, isotype antibody (Rtx: rituximab), or isotype AbLec (R7: rituximab x Siglec-7) controls alone or in combination with a CD47 antagonist antibody. Data are mean ± s.d. of replicates from n = 5 human donors.

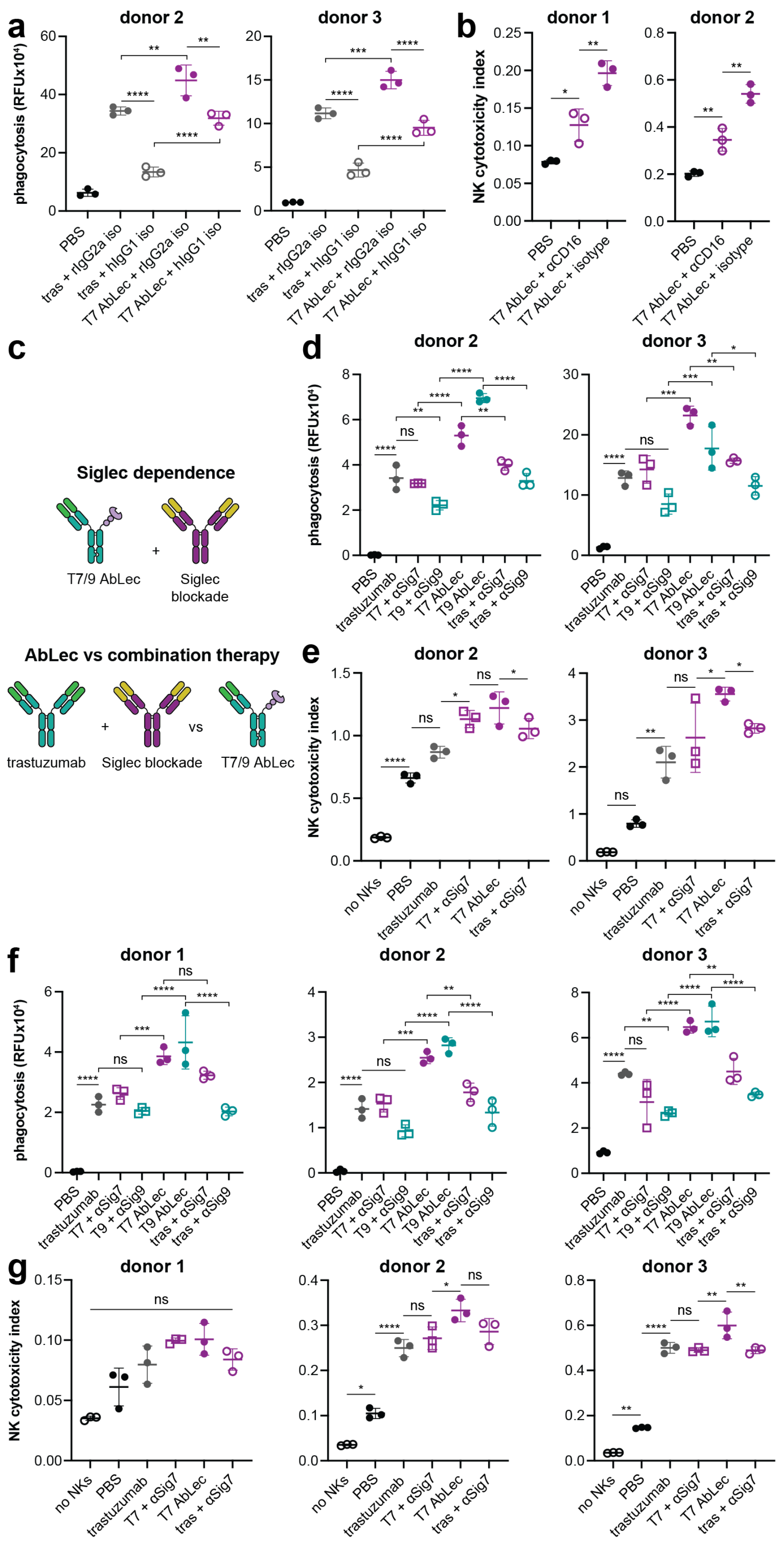
a
donor 2
donor 3
phagocytosis (RFUx10⁴)
PBS
tras + rIgG2a iso
tras + hIgG1 iso
T7 AbLec + rIgG2a iso
T7 AbLec + hIgG1 iso
b
donor 1
donor 2
NK cytotoxicity index
PBS
T7 AbLec + αCD16
T7 AbLec + isotype
c
Siglec dependence
T7/9 AbLec
+
Siglec blockade
AbLec vs combination therapy
trastuzumab
Siglec blockade
vs
T7/9 AbLec
d
donor 2
donor 3
phagocytosis (RFUx10⁴)
PBS
trastuzumab
T7 + αSig7
T9 + αSig9
T7 AbLec
T9 AbLec
tras + αSig7
tras + αSig9
e
donor 2
donor 3
NK cytotoxicity index
no NKs
PBS
trastuzumab
T7 + αSig7
T7 AbLec
tras + αSig7
f
donor 1
donor 2
donor 3
phagocytosis (RFUx10⁴)
g
donor 1
donor 2
donor 3
NK cytotoxicity index
ns

**Extended Data Figure 7. AbLecs enhance ADCC and ADCP via glyco-immune checkpoint blockade.**

(**a**) Primary human macrophages were pre-treated with non-targeting antibodies to block human FcRs (hIgG1 isotype) or non-FcR binding controls (rIgG2a isotype) prior to co-culture with SK-BR-3 cells and treatment with trastuzumab (tras) or T7 AbLec. Data are mean ± s.d. of n = 3 biological replicates.

(**b**) Primary human NK cells were pre-treated with an FcγRIII/CD16-blocking antibody (clone 3G8; αCD16) or an isotype control antibody prior to co-culture with SK-BR-3 cells and treatment with T7 AbLec. Data are mean ± s.d. of n = 3 biological replicates.

(**c**) Siglec-7 and -9 blocking antibodies were used to assess whether AbLec-mediated immune enhancement was dependent on the targeted Siglec immune checkpoint (**top**). We also compared combination of trastuzumab with Siglec blocking antibodies (αSig7 or αSig9) to T7 or T9 AbLec treatment alone in phagocytosis assays (**bottom**).

(**d, f**) Macrophage ADCP of SK-BR-3 (**d**) or HER2+ K562 (**f**) cells induced by trastuzumab or T7/9 AbLecs in the presence or absence of Siglec-7/9 blocking antibodies (αSig7 or αSig9, respectively). Data are mean ± s.d. of n = 3 biological replicates.

(**e, g**) NK cell ADCC of SK-BR-3 (**e**) or HER2+ K562 (**g**) cells induced by trastuzumab or T7 AbLec in the presence or absence of a Siglec-7 antagonist antibody (αSig7). Data are mean ± s.d. of n = 3 biological replicates.

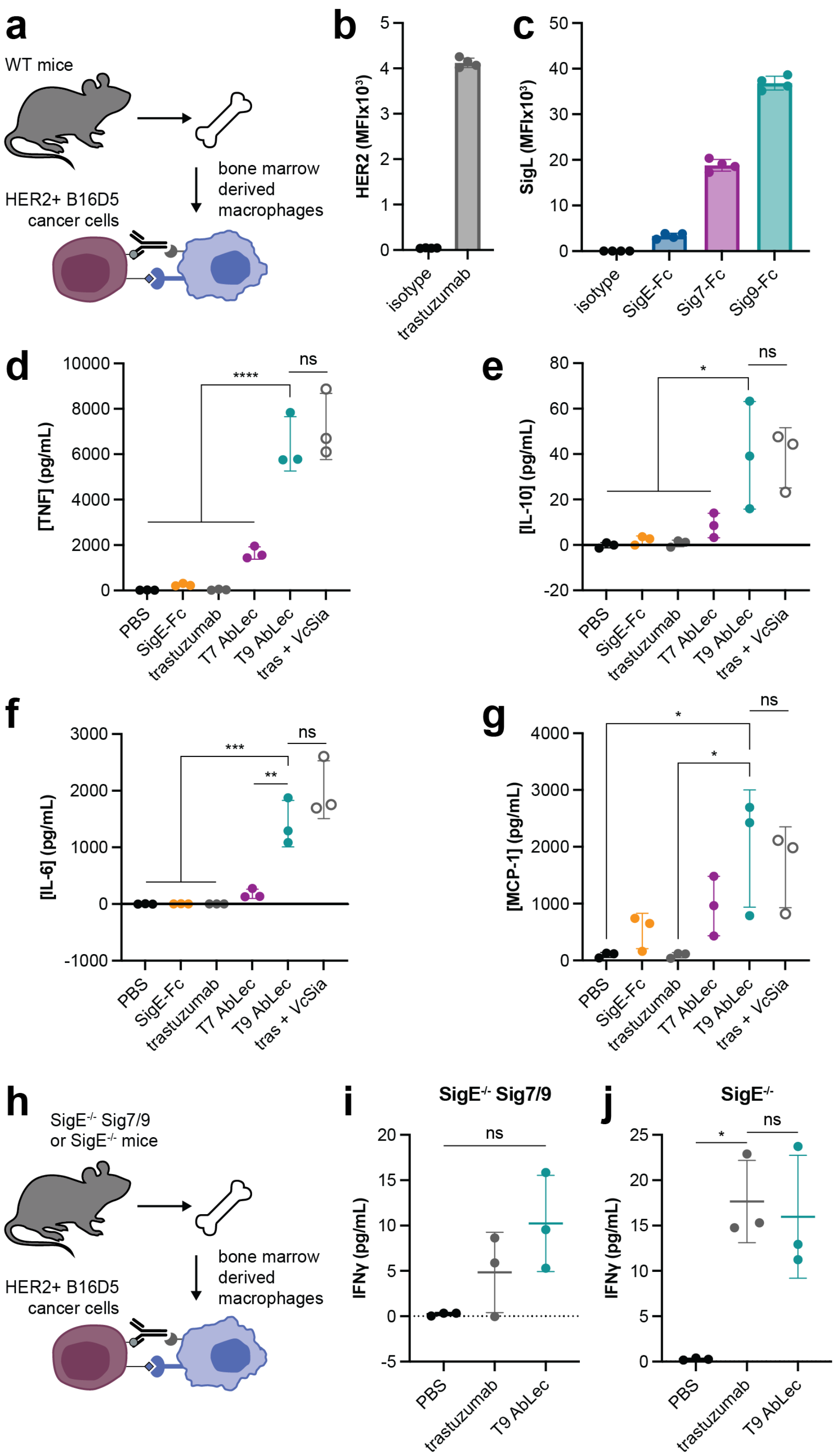


**Extended Data Figure 8. AbLecs activate murine BMDMs in a Siglec-dependent manner.**

(**a**) We tested the capacity of AbLecs to activate murine BMDMs from WT mice.

(**b, c**) Expression of HER2 (**b**) and ligands for Siglec-E, Siglec-7, and Siglec-9 on HER2+ B16D5 cells measured via flow cytometry. Data are mean ± s.d. from n = 4 biological replicates.

(**d-g**) Activation of BMDMs measured by TNF (**d**), IL-10 (**e**), IL-6 (**f**), and MCP-6 (**g**) production in BMDM/HER2+ B16D5 co-cultures treated with trastuzumab, AbLecs, or Siglec-E decoy receptor. Data are mean ± s.d. from n = 3 biological replicates.

(**h**) We tested the Siglec-dependence of AbLec activity in assays with BMDMs from Siglec-E knockout mice ($SigE^{-/-}$) and mice humanized for Siglec-7/9 receptors ($SigE^{-/-}$ Sig7/9).

(**i**) Activation of $SigE^{-/-}$ Sig7/9 BMDMs measured by IFNγ production in BMDM/HER2+ B16D5 co-cultures treated with trastuzumab or T9 AbLec. Data are mean ± s.d. from n = 3 biological replicates.

(**j**) Activation of $SigE^{-/-}$ BMDMs measured by IFNγ production in BMDM/HER2+ B16D5 co-cultures treated with trastuzumab or T9 AbLec. Data are mean ± s.d. from n = 3 biological replicates.

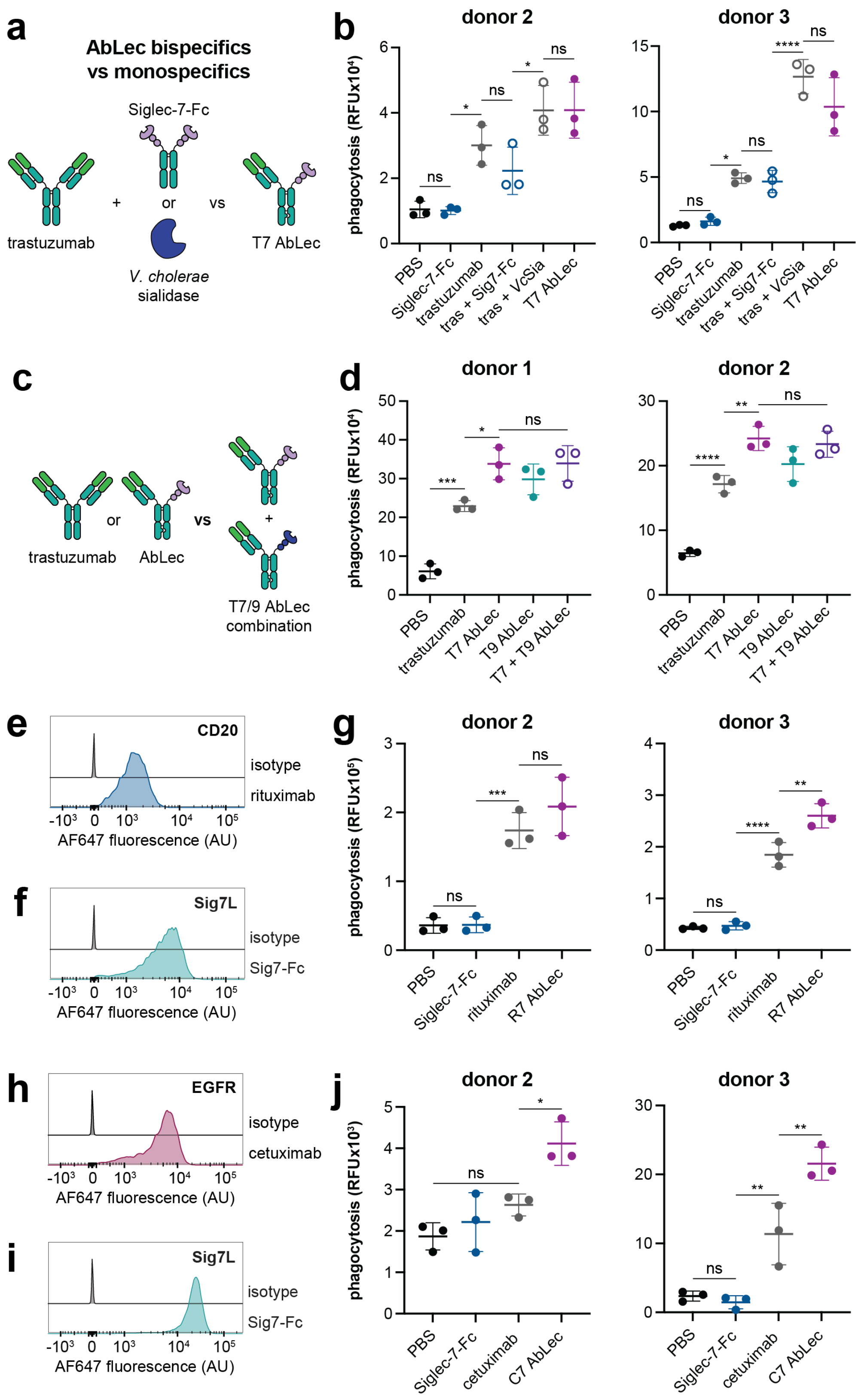


**Extended Data Figure 9. The chimeric AbLec architecture enables targeting of diverse cancers.**

(**a**) We compared AbLec treatment to combination of trastuzumab and a decoy receptor to test whether the chimeric AbLec architecture was required for efficacy.

(**b**) ADCP of HER2+ K562 cells by primary human macrophages *in vitro* in co-cultures treated with Siglec-7-Fc, trastuzumab, T7 AbLec, or combinations of trastuzumab with Siglec-7-Fc or *V. cholerae* sialidase (*Vc*Sia). Data are mean ± s.d. of n = 3 biological replicates.

(**c**) We asked whether combination of two AbLecs targeting distinct glyco-immune checkpoints could further enhance immune cell activation.

(**d**) ADCP of SK-BR-3 cells by primary human macrophages in co-cultures treated with equimolar concentrations of trastuzumab, T7 AbLec, T9 AbLec, or combination of T7 and T9 AbLecs. Data are mean ± s.d. of n = 3 biological replicates.

(**e, f**) Ramos cells expressed CD20 (**e**) and Sig7L (**f**) by flow cytometry. Histograms are representative of n = 3 biological replicates.

(**g**) ADCP of Ramos cells by primary human macrophages in co-cultures treated with R7 AbLec, rituximab, or Siglec-7-Fc. Data are mean ± s.d. of n = 3 biological replicates.

(**h, i**) EGFR+ K562 cells expressed EGFR (**h**) and Sig7L (**i**) by flow cytometry. Histograms are representative of n = 3 biological replicates.

(**j**) ADCP of EGFR+ K562 cells by primary human macrophages in co-cultures treated with C7 AbLec, cetuximab, or Siglec-7-Fc. Data are mean ± s.d. of n = 3 biological replicates.

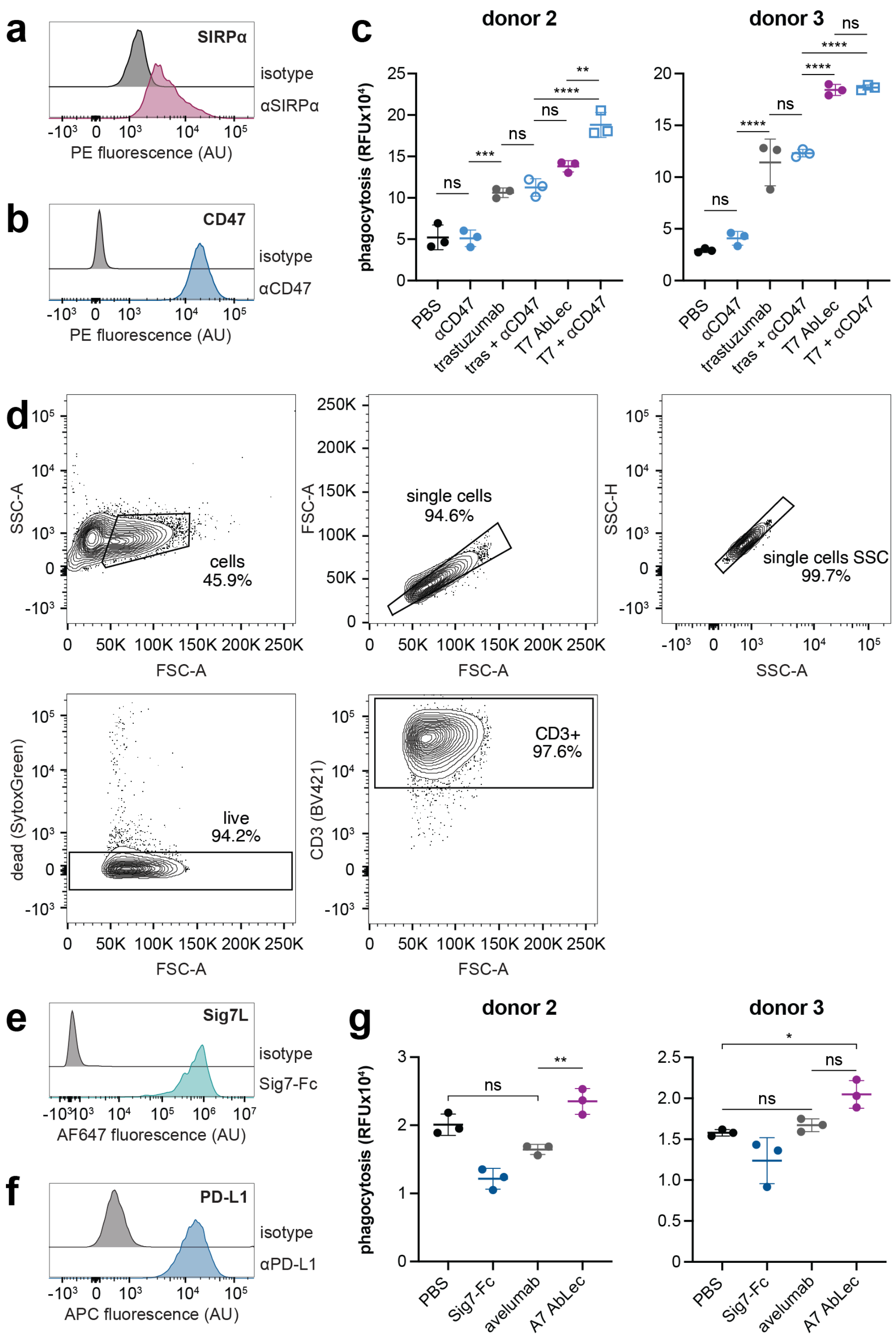


**Extended Data Figure 10. AbLecs synergize with blockade of established immune checkpoints.**

(**a**) Primary human macrophages used in *in vitro* phagocytosis assays express SIRPα by flow cytometry. Histogram is representative of biological replicates from n = 2 donors.

(**b**) SK-BR-3 cells express CD47 by flow cytometry. Histogram is representative of n = 2 biological replicates.

(**c**) ADCP of SK-BR-3 cells by primary human macrophages in co-cultures treated with αCD47 antagonist antibody, trastuzumab, and T7 AbLec alone and in various combinations. Data are mean ± s.d. of n = 3 biological replicates.

(**d**) Human T cells were defined as CD3+ by flow cytometry.

(**e, f**) MDA-MB-231 cells express Sig7L (**e**) and PD-L1 (**f**) by flow cytometry. Histograms are representative of n = 2 biological replicates.

(**g**) ADCP of SK-BR-3 cells by primary human macrophages in co-cultures treated with A7 AbLec, avelumab (αPD-L1), or Siglec-7-Fc. Data are mean ± s.d. of n = 3 biological replicates.
